# Comparison of nuisance function construction strategies for double machine learning causal inference in single-cell transcriptomics: shared unsupervised deep learning does not require cross-fitting

**DOI:** 10.64898/2026.07.22.739971

**Authors:** Wei Ye, Xinyu Jiang, Feng(Ben) Shen

## Abstract

Inferring “whether a change in the expression of a given gene causally affects the disease state” from observational single-cell transcriptomic data is one of the central problems in single-cell biology. The difficulty lies in confounding: cell state, batch, cell cycle, and the co-expression of other genes may all simultaneously influence the target gene (treatment variable T) and the disease label (outcome variable Y), so that naive correlation analysis cannot distinguish causation from covariation. Double machine learning (DML), via orthogonal scores and cross-fitting, allows machine learning to estimate high-dimensional nuisance functions, thereby addressing the causal inference problem in high-dimensional data. Constrained by computational resources, this study takes a small-sample dataset with p≈n (2,120 cells, 1,999 background genes) as the experimental testbed and systematically compares three nuisance function construction strategies under this critical condition; strategies for the n≫p regime are then addressed by theoretical argument. Using systemic lupus erythematosus (SLE) peripheral blood memory B cells (GSE189050, 2,120 cells), we construct a **three nuisance function construction strategies x (in-sample / cross-fitting)** 2×3 factorial experiment and compare them in terms of resolution, biological plausibility, stability, and deconfounding ability in causal effect estimation. The three strategies are: (S1) direct linear nuisance regression on the high-dimensional background genes without dimensionality reduction; (S2) learning a shared low-dimensional representation with an unsupervised autoencoder, with the treatment and outcome residuals sharing that representation; (S3) fitting the treatment and outcome with two independent deep networks.

The results yield a clear three-part pattern with mechanistic meaning: **(1) both modes of S1 fail; (2) S2 achieves the optimum in-sample and requires no cross-fitting; (3) S3 is “rescued” under cross-fitting and becomes effective.** We explain this pattern starting from the convergence rate condition of DML (Sec.5 Theoretical Foundations) and give the applicability boundaries of the two strategies: **S2 (shared unsupervised deep learning) achieves estimation quality comparable to standard cross-fitted DML at a tiny fraction of the computational cost, making it a feasible scheme for large-scale screening; S3 (dual networks + cross-fitting), as the standard DML recipe, can serve as a broad-spectrum reference control for S2 results.**

## 1. Introduction

### 1.1 Background

Inferring “whether a change in the expression of a given gene causally affects the disease state” from observational transcriptomic data is one of the central problems in single-cell biology. The difficulty lies in confounding: cell state, batch, cell cycle, and the co-expression of other genes may all simultaneously influence the target gene (treatment variable T) and the disease label (outcome variable Y), so that naive correlation analysis cannot distinguish causation from covariation.

Semiparametric methods for causal inference have opened a new path for this problem. Double machine learning (DML; ^[1]^) decomposes the estimation error of the nuisance functions and the causal parameter via the Neyman orthogonal score, and has become the standard framework for causal effect estimation from high-dimensional observational data.

In the single-cell setting, a series of causal inference and perturbation-effect prediction methods have recently emerged: CINEMA-OT identifies single-cell experimental perturbation effects via optimal transport ^[2]^; generative models such as scCausalVI disentangle perturbation responses through causality-aware representations ^[3]^; causal graph neural networks such as GRACE infer gene regulatory networks from expression profiles ^[4]^; SINCERITIES infers gene regulatory relationships from time-stamped single-cell expression data ^[5]^; Perturb-seq combines CRISPR screening with single-cell RNA sequencing to enable large-scale perturbation-effect profiling ^[6]^; RENGE infers gene regulatory networks from time-series CRISPR perturbation data ^[7]^; Scribe infers causal gene regulatory networks from coupled single-cell expression dynamics ^[8]^; and multiple machine learning models have been validated for perturbation-effect prediction ^[9]^; benchmarking studies show that Single-cell perturbation technologies enable systematic investigation of gene functions and regulatory networks with single-cell resolution ^[10]; [11]^.yet **the relative performance of different nuisance function construction strategies in single-cell DML lacks systematic comparison.**

The core insight of DML is that, as long as the estimation errors of the two nuisance functions satisfy the **product rate condition**

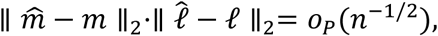

i.e., each achieves *o*_*P*_(*n*^−1/4^), then the causal parameter 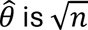-consistent and asymptotically normal—the first-order bias from nuisance function estimation is canceled by the orthogonal score. This means that DML is **means-neutral** with respect to the choice of nuisance function: any method that can estimate the nuisance functions at an *o*_*P*_(*n*^−1/4^) rate—whether a regularized linear method such as Lasso, an ensemble method such as a random forest, or a deep neural network ^[12]; [13]^—can in principle serve as the fitter. But means-neutrality is not means-irrelevance: in a concrete application, which method can actually attain this rate depends on how well the data characteristics match the method’s structural assumptions (Sec.5.1–5.2). The complete derivation of this rate condition is given in Sec.5.1.

### 1.2 Problem: in single-cell DML, how to construct nuisance functions to satisfy the product rate condition becomes the key

This study takes a small-sample dataset with p≈n (2,120 cells, 1,999 background genes) as the experimental testbed and systematically compares three nuisance function construction strategies under this condition. This choice rests on two considerations: first, the computational constraint—each S3cf estimate requires 2 deep networks x 5-fold training, and cross-fitting at the scale of thousands of genes across the full transcriptome is infeasible under current computational resources, so the small sample makes a complete 2×3 factorial comparison possible; second, the bias-dominated failure mechanism under n≫p, although more aligned with real application scenarios, is already established independently by the DML rate condition and Stone’s nonparametric theory (Sec.5.1–5.2, see the n≫p extrapolation in Sec.5.2) and need not rely on experimental extrapolation.

Low-dimensional representations provide a powerful tool for constructing nuisance functions. Unsupervised dimensionality-reduction methods such as the autoencoder can map high-dimensional transcriptomes onto a low-dimensional manifold representation, preserving biological variation while removing technical noise^[14]^, scVI generative dimensionality reduction ^[15]^, DCA denoising autoencoder^[16]^, scTFBridge disentangled generative model^[17]^, interpretation and inference of single-cell perturbation). In addition, causal generative models such as GRouNdGAN demonstrate the ability to simulate single-cell data starting from gene regulatory networks ^[18]^, while methods based on time-series causal inference ^[19]^ and counterfactual deep-learning frameworks ^[20]^ further enrich the toolbox for single-cell causal inference. Such low-dimensional representations are naturally suited as inputs to DML, but two diametrically opposed usages: one is **sharing a single dimensionality reduction** (treatment and outcome share the same representation), and the other is **learning independent representations** for the treatment and outcome separately. The empirical performance and theoretical properties of the two approaches lack direct comparison. Therefore, **the relative performance of different nuisance function construction strategies in single-cell DML warrants systematic comparison**—this is precisely the starting point of the present study.

### 1.3 Three nuisance function construction strategies and the design of this study

Centered on “how to construct the nuisance functions,” we distill three representative strategies that span the design space from the simplest to the most complex:

- **S1 direct DML (no dimensionality reduction)**: fit the two nuisance functions directly on the high-dimensional background X with linear models.
- **S2 shared unsupervised deep learning + DML**: first compress X into a low-dimensional representation Z with an unsupervised autoencoder, then fit the treatment and outcome residuals on the **same Z** with linear regression. “Shared” means the two nuisance functions reuse the same representation.
- **S3 two independent deep networks + DML**: fit *E*[*T*|*X*] and *E*[*Y*|*X*] separately with two independent supervised networks. This is the canonical implementation of standard DML.

For each strategy we further cross two fitting modes: **in-sample**—train on the full sample, predict on the full sample; **cross-fitting**—5-fold, train on the other 4 folds for each fold and predict on the held-out fold, then concatenate the out-of-sample residuals (the standard DML recipe). This yields **2×3 = 6 combinations**, enabling horizontal (representation method), vertical (cross-fitting effect), and interaction comparisons. The data-flow differences among the three strategies in nuisance function construction are shown in Figure 1.

**Figure 1.**
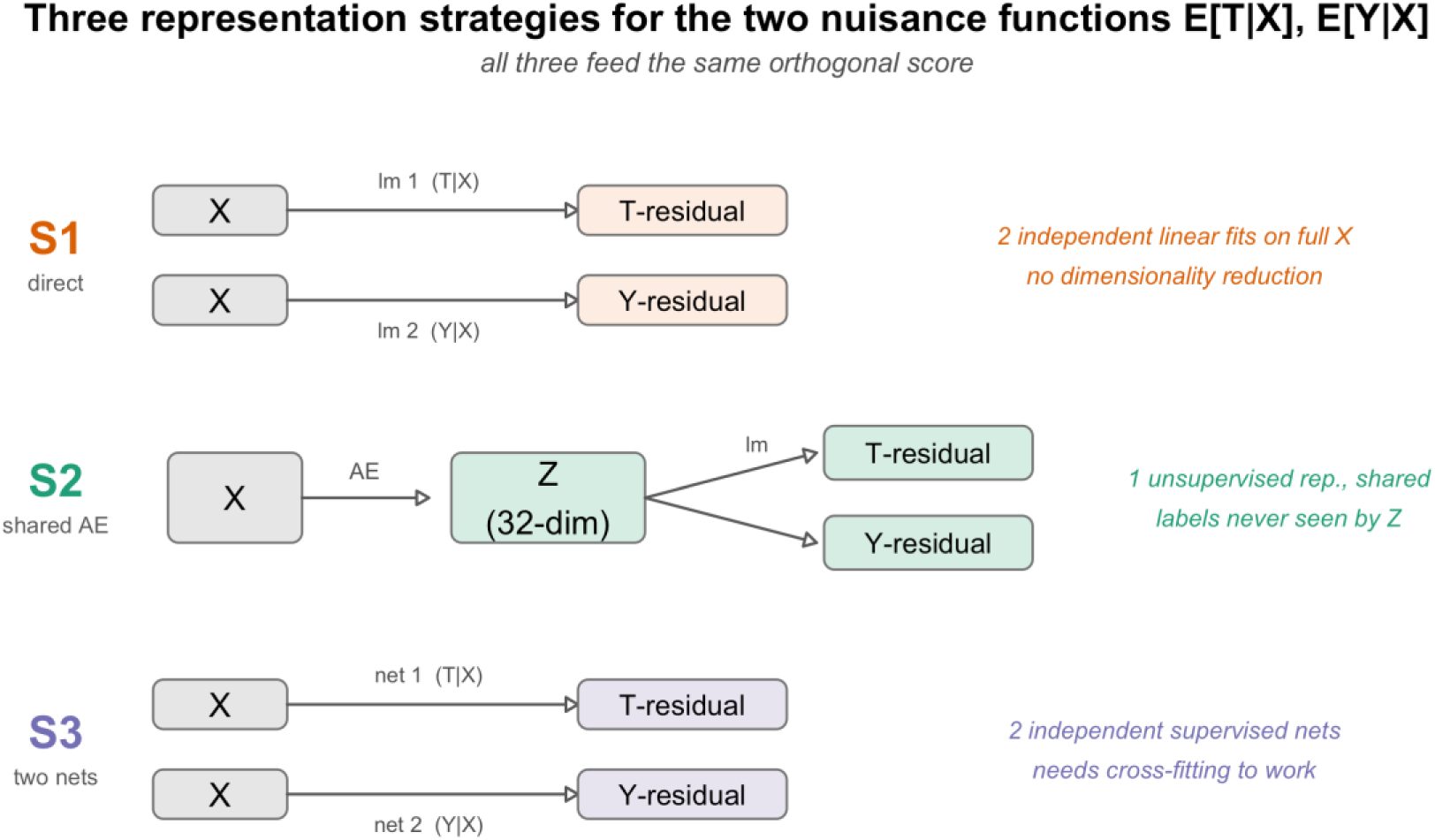
Architecture of the three representation strategies: S1 directly fits two independent linear regressions on high-dimensional X for E[T|X] and E[Y|X] (no dimensionality reduction); S2 uses one unsupervised autoencoder to compress X into a 32-dimensional shared representation Z, with both residuals sharing Z for linear regression; S3 uses two independent supervised networks to fit E[T|X] and E[Y|X] separately. All three share the same orthogonal score at the final step.

## 2. Methods

### 2.1 Data and preprocessing

The data were taken from GSE189050^[21]^ (SLE peripheral blood mononuclear cells). Cells annotated as **Memory B cells** with disease group **SLE ACT (active)** or **Control** were selected, yielding **2,120 cells** (634 cases, 1,486 controls). Raw counts from the RNA assay were standard log-normalized (LogNormalize, scale factor = 1e4).

The top 2,000 highly variable genes formed the analysis gene pool.

### 2.2 Treatment, outcome, and background (leave-one-out)

- **Outcome Y**: binary disease status (SLE ACT = 1, Control = 0).
- **Target genes**: 30 genes randomly drawn from the highly variable gene pool (fixed seed 20260714), each serving in turn as the treatment variable T for estimating its causal effect.
- **Background X (leave-one-out)**: when estimating gene g, the background = all highly variable genes **except g** (p = 1,999); the remaining 29 target genes whose turn has not yet come **remain in the background** as confounders and are not permanently removed.

### 2.3 Implementation of the three strategies (key: align the estimators as far as possible, leaving only the difference under investigation)

**S1**: directly fit lm(T ∼ X₋g) and glm(Y ∼ X₋g, binomial) on the high-dimensional background X₋g, take the residuals, and compute the orthogonal score. **S1 and S2 use exactly the same DML estimator (linear nuisance regression + orthogonal score); the only difference is that S1 does not reduce dimension**—so if S1 fails, it can only be attributed to “no dimensionality reduction,” giving a clean control. (Note: S1 in this paper refers specifically to **unregularized** direct linear DML; the status of regularized methods such as Lasso is discussed in Sec.6.1.)

**S2**: for each gene g, train an autoencoder on the background X₋g and take the 32-dimensional bottleneck representation Z_g; both the treatment and outcome residuals are linearly fitted on the **same Z_g** with lm/glm. The autoencoder has an 8-layer symmetric structure (encoder 4096->2048->1024->512->256->128->64->32, decoder symmetric), Adam optimizer, learning rate 5e-4, 35 epochs, batch 256.

**S3**: for each gene g, use two **independent** supervised networks to predict T and Y separately. The backbone of both networks is **identical** to the S2 encoder (4096->…->32, same learning rate/epochs/batch). Thus **S2 and S3 have identical network capacity and hyperparameters**; the only essential difference is that S2 is unsupervised reconstruction with the two residuals sharing a representation, whereas S3 uses two supervised networks each learning its own representation.

### 2.4 Cross-fitting

In the cross-fitting mode, the 2,120 cells are randomly split into 5 folds. For fold k, the nuisance functions (S2’s autoencoder and linear head, S3’s two networks) are trained on the other 4 folds, and **out-of-sample** prediction is made on the held-out fold k; the 5 folds are concatenated to obtain the out-of-sample residual for each cell, and the orthogonal score is then computed. **Both S2’s autoencoder and S3’s networks strictly follow “train within the fold, predict on the held-out fold”**—the representation never touches the held-out fold, consistent with the cross-fitting specification of Chernozhukov et al. (2018); S2cf and S3cf are fully comparable on this point.

### 2.5 Evaluation metrics

1. **Inter-gene variance (resolution)**: the variance of θ̂ across the 30 genes; the larger, the better the method distinguishes the strength of effects among genes.
2. **Intra-gene estimation variance (stability)**: bootstrap-resample the sample, re-estimate θ̂, and take the variance of repeated estimates of the same gene; the smaller, the more stable. For the three in-sample strategies we use N = 6 bootstraps; the stability of S3cf is assessed separately (see Sec.3.4).
3. **Rank correlation (biological plausibility)**: the rank correlation between |θ̂| and the marginal |gene–disease Spearman correlation|; the higher, the more the ranking of causal-effect strength accords with the known marginal association.
4. **Residual confounding diagnostic**: the mean absolute correlation of the residuals T_res, Y_res with the background X; it reflects whether the nuisance functions have fully absorbed the confounding information. This metric is **one-directional**: provided the residuals have not been crushed by overfitting, higher correlation means less adequate absorption of confounding (informative); but low correlation does not necessarily mean adequate absorption—when the model overfits and interpolates, the residuals are compressed to numerical zero, feigning an extremely low correlation (see Sec.3.6). We therefore use it only as an auxiliary diagnostic, not as a primary criterion.
5. **Qualitative observations**: the θ̂ sign-consistency matrix; the shrinkage-factor regression relative to S2 (in-sample).

## 3. Results

### 3.1 Magnitude of θ̂: who collapses

The typical magnitudes of θ̂ across the combinations are distinguishable at a glance (median |θ̂|):

| Strategy | In-sample | Cross-fit |
| --- | --- | --- |
| **S1 direct** | $\sim 10^{-12}$ (collapsed) | $\sim 10^{-3}$ (slight recovery, still weak) |
| **S2 shared deep learning** | $\sim 10^{-2}$ (healthy) | $\sim 10^{-2}$ (healthy) |
| **S3 dual independent** | $\sim 10^{-6}$ (collapsed) | $\sim 10^{-2}$ (rescued) |

S1’s in-sample θ̂ falls in the 10⁻¹² range, which is essentially numerical zero-high-dimensional unregularized linear regression interpolates the residuals to near 0, and the orthogonal score θ̂ = Σ(*T̃Ỹ*)/Σ(T̃²) degenerates to 0/0. S3 in-sample falls at 10⁻⁶, likewise a result of the residuals carrying almost no information after representation collapse. Only S2 yields θ̂ at the 10⁻² level in-sample-meaningful causal-effect estimates. The magnitudes of θ̂ across the combinations are shown in Figure 2.

**Figure 2.**
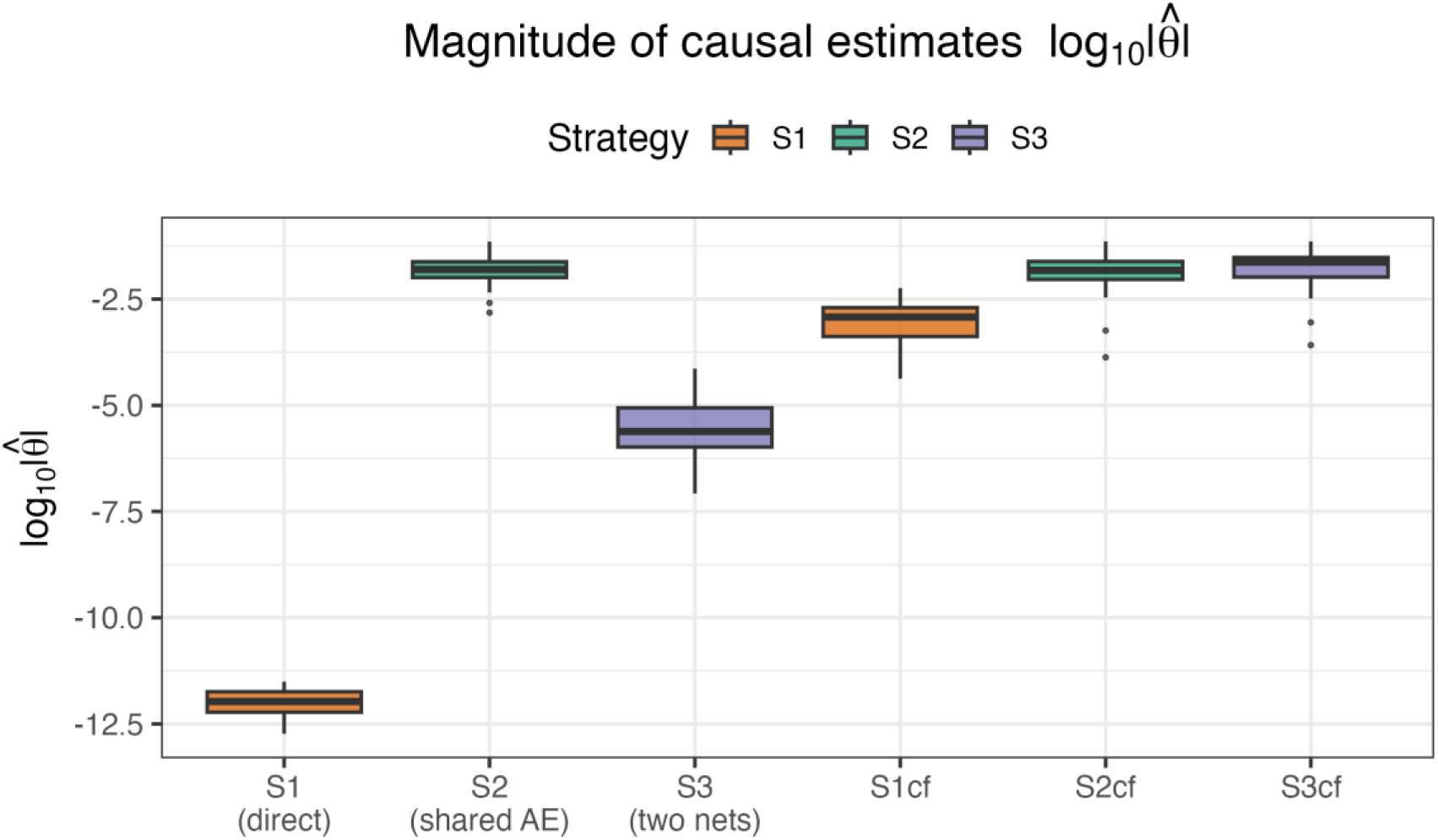
Magnitude of causal effect estimates (log₁₀|θ̂|) across the six combinations. S1 and S3 in-sample collapse to near numerical zero.

### 3.2 Resolution and biological plausibility

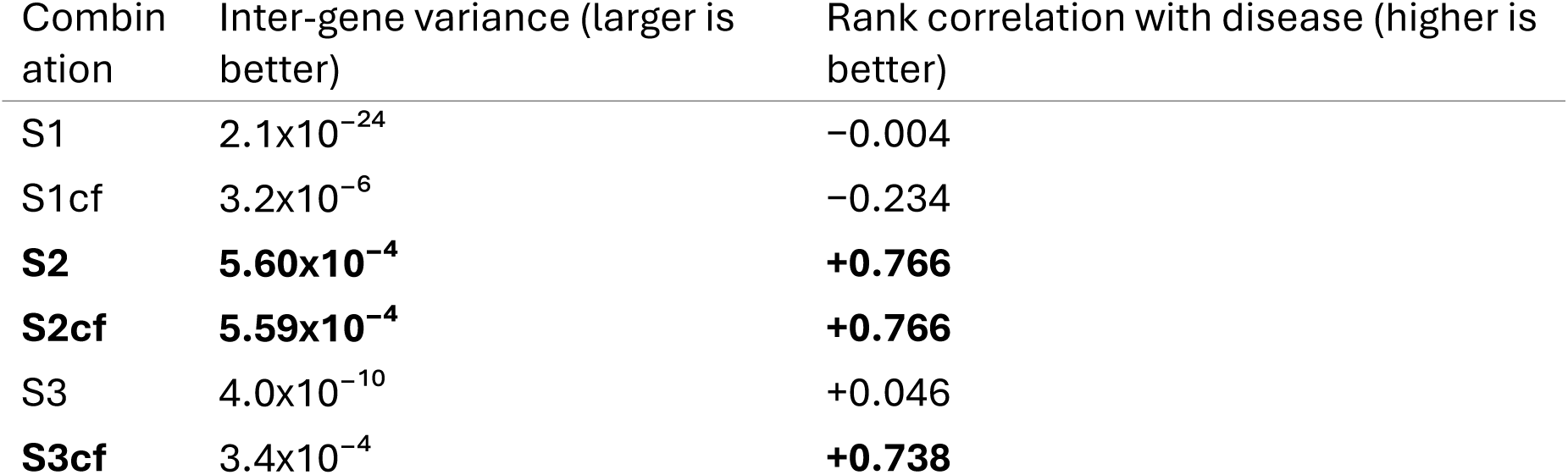

| Combination | Inter-gene variance (larger is better) | Rank correlation with disease (higher is better) |
| --- | --- | --- |
| S1 | $2.1 \times 10^{-24}$ | -0.004 |
| S1cf | $3.2 \times 10^{-6}$ | -0.234 |
| **S2** | **$5.60 \times 10^{-4}$** | **+0.766** |
| **S2cf** | **$5.59 \times 10^{-4}$** | **+0.766** |
| S3 | $4.0 \times 10^{-10}$ | +0.046 |
| **S3cf** | **$3.4 \times 10^{-4}$** | **+0.738** |

Three key points:

1. **S2 ≈ S2cf**: in both resolution (5.60 vs 5.59 x10⁻⁴) and biological rank correlation (0.766 vs 0.766), the in-sample and cross-fitted versions of S2 are virtually identical-cross-fitting brings no change whatsoever to S2.
2. **S3 is rescued by cross-fitting**: the in-sample rank correlation of S3 is only 0.046 (biologically meaningless); after cross-fitting, S3cf’s rank correlation jumps to 0.738 and its resolution improves by about six orders of magnitude (10⁻¹⁰ -> 10⁻⁴), recovering to a level close to S2.
3. **S1 cannot be rescued**: S1cf’s rank correlation is −0.234 (not statistically significant at n = 30, at the same noise level as S1 in-sample’s −0.004); although resolution rises from 10⁻²⁴ to 10⁻⁶, it is still about two orders of magnitude below S2-cross-fitting cannot remedy the structural defect of “no dimensionality reduction.” The resolution and rank correlation across the combinations are shown in Figures 3–4, and the differential effect of cross-fitting on the three strategies is shown in Figure 5.

**Figure 3.**
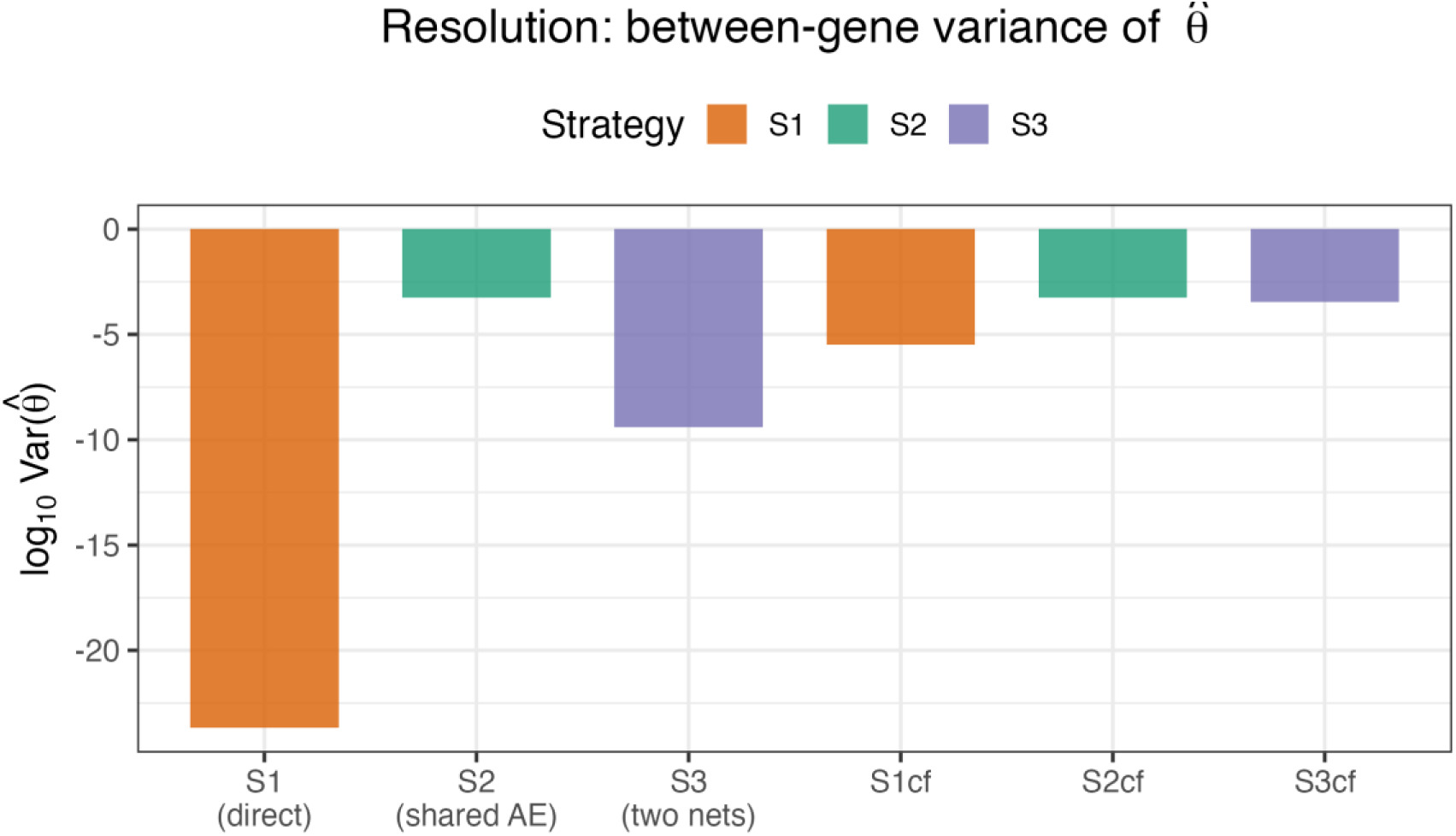
Inter-gene variance (resolution, log scale).

**Figure 4.**
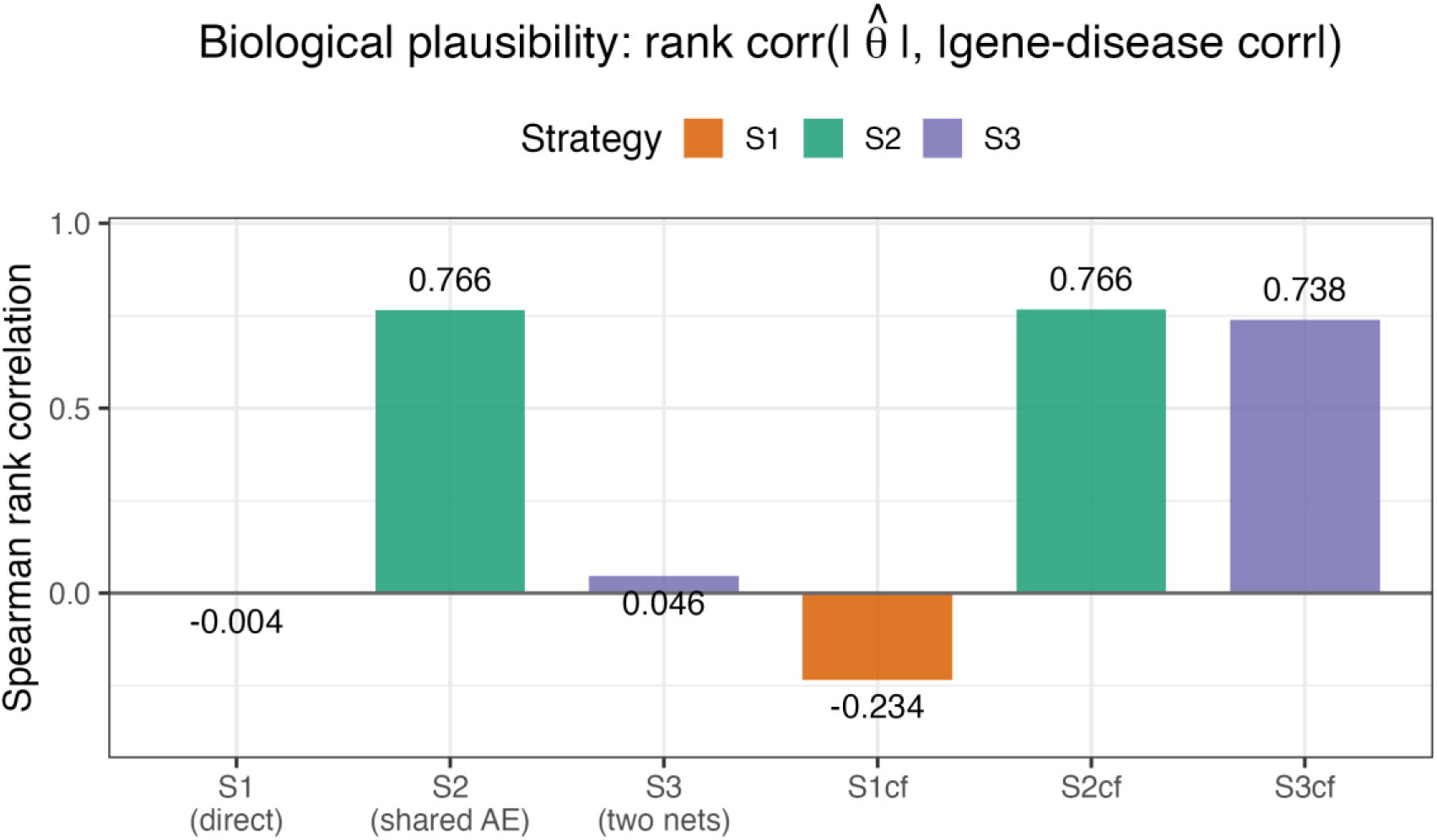
Spearman rank correlation between |θ̂| and |gene–disease marginal correlation| (biological plausibility, core figure).

**Figure 5.**
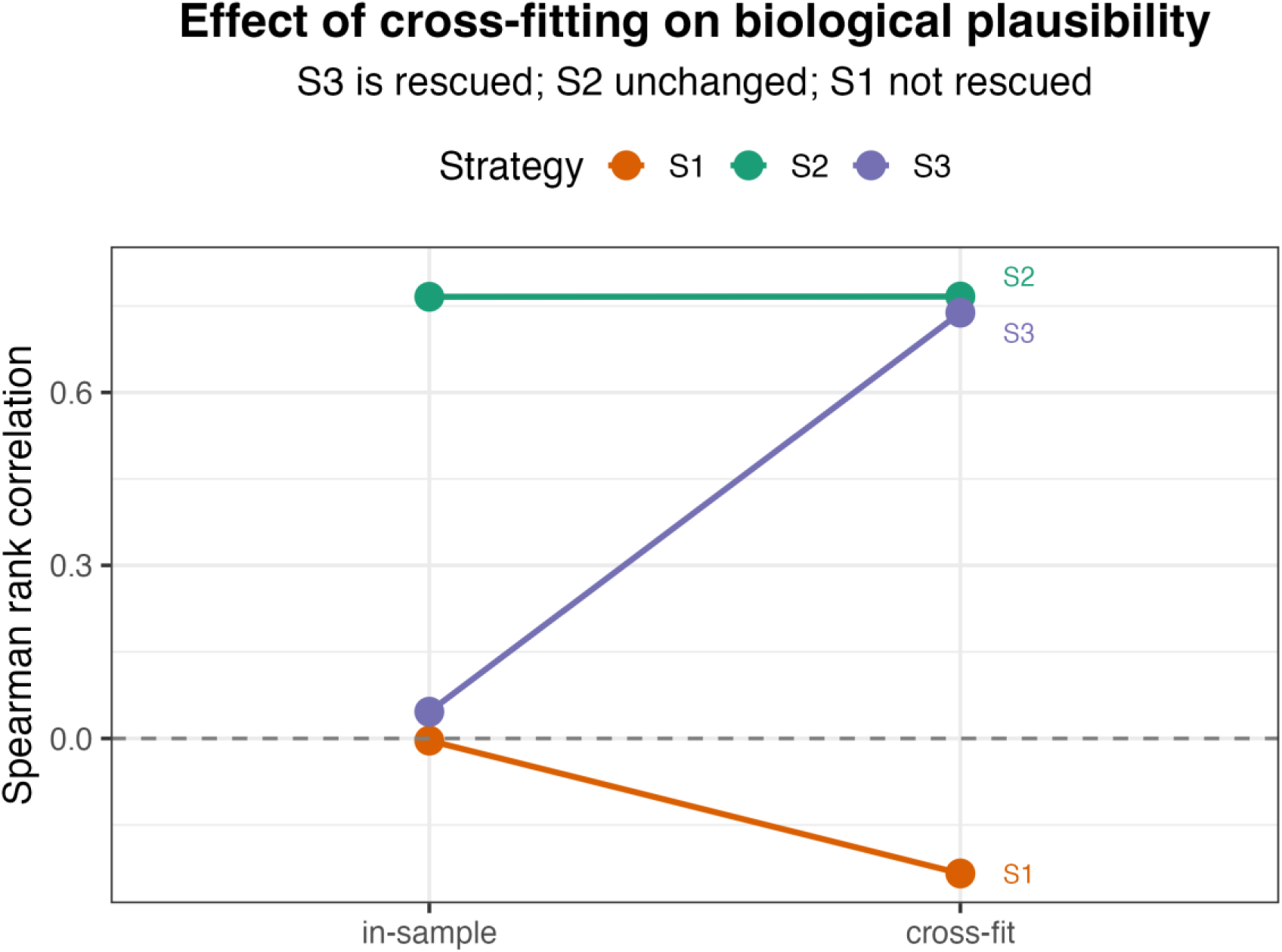
Effect of cross-fitting on biological plausibility: S3 rescued, S2 unchanged, S1 not rescued.

Plotting |θ̂| against the |gene–disease marginal correlation| gene by gene for the six combinations (Figure 6) makes this pattern visible directly: the scatter of S2, S2cf, and S3cf aligns along a positive trend (rank correlation above 0.7), whereas that of S1, S1cf, and S3 is a nearly structureless noise cloud.

**Figure 6.**
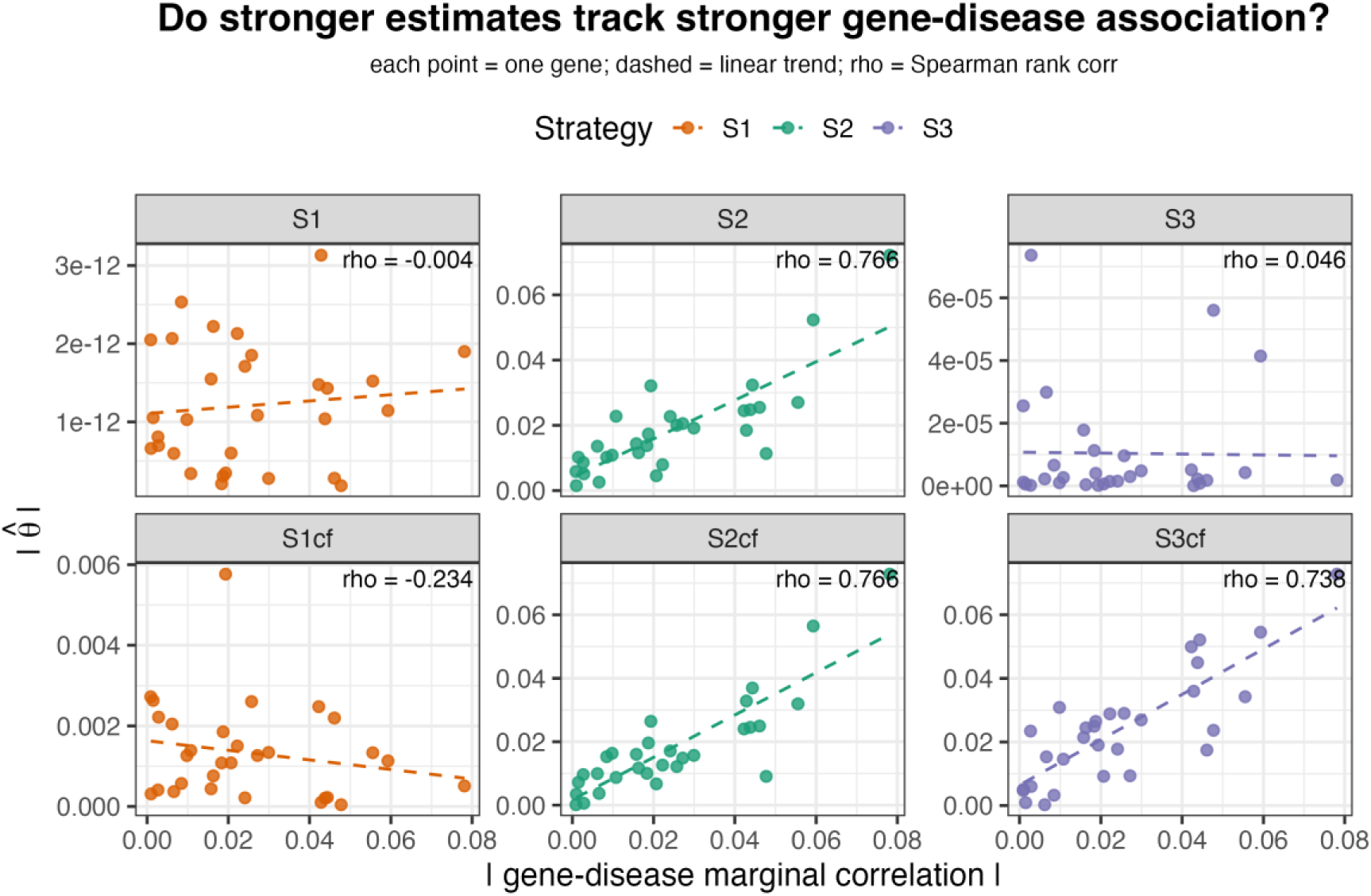
Gene-wise scatter plots of |θ̂| against |gene–disease marginal correlation| for all six combinations (each point is one gene; dashed line shows linear trend; ρ is Spearman rank correlation). S2/S2cf/S3cf exhibit a positive trend, while S1/S1cf/S3 form noise clouds.

### 3.3 Stability (intra-gene bootstrap variance)

The mean intra-gene variance for the three in-sample strategies (N = 6 bootstraps):

| Strategy (in-sample) | Intra-gene variance<br>(smaller is more<br>stable) | Interpretation |
| --- | --- | --- |
| S1 | 8.78 | Extremely unstable: high-dimensional lm fluctuates violently across bootstrap samples |
| S2 | $2.7 \times 10^{-4}$ | Stable |
| S3 | $1.3 \times 10^{-10}$ | False stability: $\theta$ has already collapsed near 0, so the variance is naturally tiny; this is not genuine stability |

The huge variance of S1 (8.78) stands in sharp contrast to the small variance of S2 (2.7×10⁻⁴). Although S3 in-sample has the smallest numerical variance, this is “stably estimating a near-zero wrong value” and should not be read as an advantage-it must be understood together with the collapse documented in Sec.3.2. The intra-gene bootstrap variance across the six combinations is compared in Figure 7.

**Figure 7.**
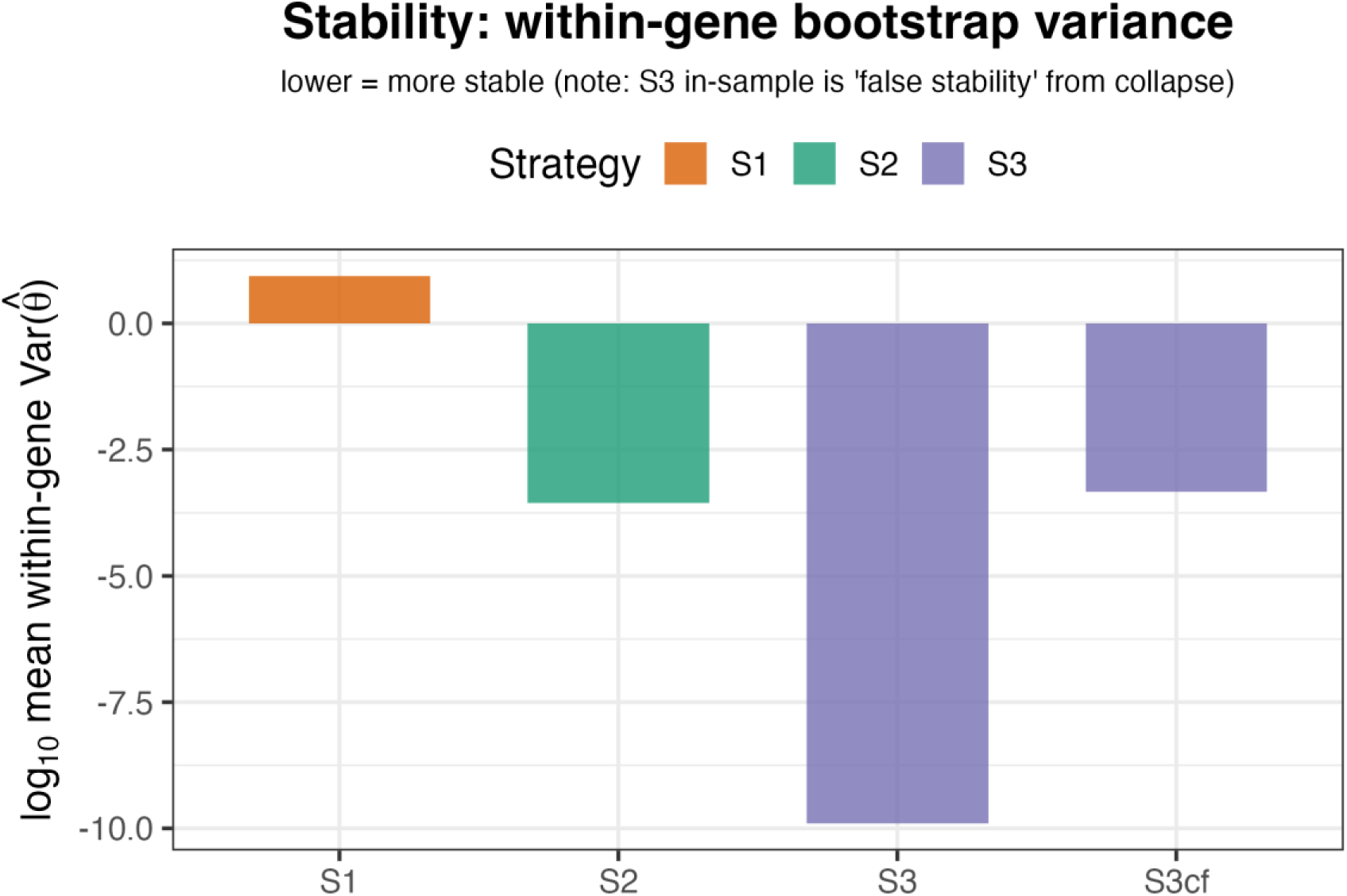
Intra-gene bootstrap variance (stability, log scale).

### 3.4 Stability of S3cf

Since S3cf has become one of the usable methods alongside S2, its stability is necessary evidence supporting “S3cf is broadly applicable.” We re-estimated θ̂ with N = 3 bootstraps on 10 genes to assess the intra-gene estimation variance of S3cf, and compared it at the order-of-magnitude level with the three in-sample strategies.

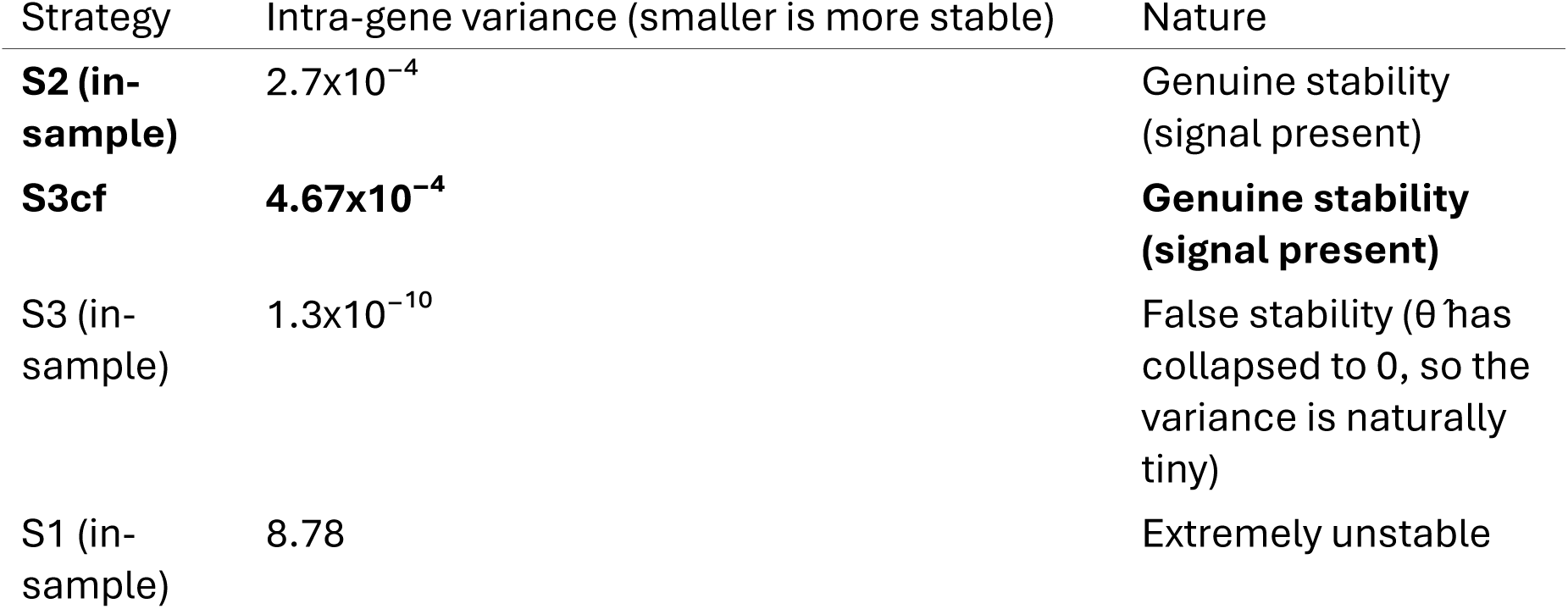

The results bear out the theoretical expectation of Sec.5.4: **the intra-gene variance of S3cf (4.67×10**⁻**⁴) is of the same order of magnitude as S2 (2.7×10**⁻**⁴)**-this is “genuine stability backed by a real signal,” fundamentally different from S3 in-sample’s “stably estimating a near-zero wrong value” (the 1.3×10⁻¹⁰ *false stability*). This confirms the usability of S3cf as the standard DML recipe; at the same time, S2’s variance is slightly smaller than S3cf’s (by about 1.7x), which, combined with S2’s greater computational economy, jointly supports the method-selection conclusion that “**S2 has a dual slight advantage in computational cost and stability, while S3cf is comparable in stability with broader theoretical assumptions**” (Sec.6.4).

### 3.5 Sign consistency and shrinkage factor

**Sign-consistency matrix** (the proportion of the 30 genes sharing the same θ̂ sign, Figure 8): the most striking result is that **the consistency of S2 and S2cf reaches 0.G33** (28 of the 30 genes share the same sign), again confirming that cross-fitting barely changes S2. By contrast, the consistency of S1 with every combination lies in 0.4–0.5 (around the random level), reflecting that its estimates are already noise.

**Figure 8.**
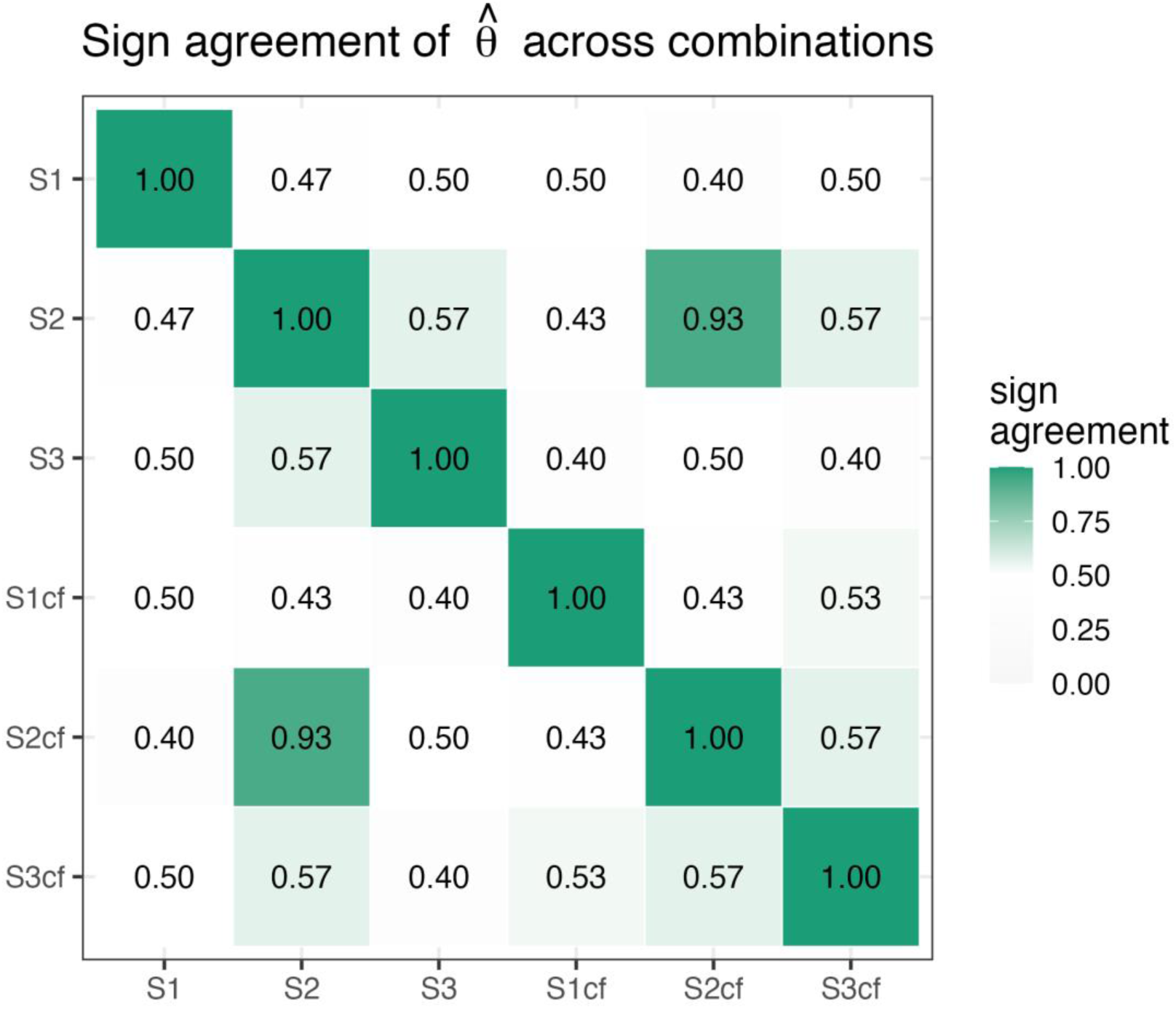
Sign consistency heatmap of θ̂ across the six combinations.

**Shrinkage factor** (the regression slope β of each combination’s θ̂ against S2 in-sample θ̂):

Here we take the S2 in-sample estimate as the reference frame for two reasons: first, S2 is the protagonist of this paper’s argument, so each method’s deviation from it most directly answers “how far the other strategies are from the central strategy”; second, the credibility of S2 is not presupposed but has been independently established first by other metrics (biological rank correlation 0.766, smallest intra-gene stability variance). It should be emphasized that the standard DML textbook recipe S3cf also converges toward S2 (β = 0.591, sign-consistent), which is tantamount to providing external endorsement for S2 from a method that is theoretically acknowledged as correct; this is therefore not circular reasoning “using the conclusion under test as the benchmark.”

| Comparison | $\beta$ | Meaning |
| --- | --- | --- |
| S2cf vs S2 | 0.975 | S2cf ≈ S2 (slope near 1) |
| S3cf vs S2 | 0.591 | S3cf is positively correlated with S2, same direction, slightly shrunk in magnitude |
| S3 vs S2 | $9.9 \times 10^{-5}$ | S3 in-sample has collapsed |
| S1 vs S2 | $5.8 \times 10^{-12}$ | S1 has completely collapsed |
| S1cf vs S2 | -0.007 | S1cf is essentially unrelated to S2 |

The β ≈ 0.975 for S2cf vs S2 and β ≈ 0.591 for S3cf vs S2 jointly show that the two usable methods after cross-fitting (S2cf, S3cf) agree with the S2 in-sample estimate in direction and are comparable in magnitude, forming a mutually corroborating set of “valid solutions”; the S1 series and S3 in-sample fall outside this set. Plotting the gene-wise θ̂ of the three usable methods as parallel coordinates (Figure 9) shows that the estimates of S2, S2cf, and S3cf are highly consistent in both sign and magnitude.

**Figure 9.**
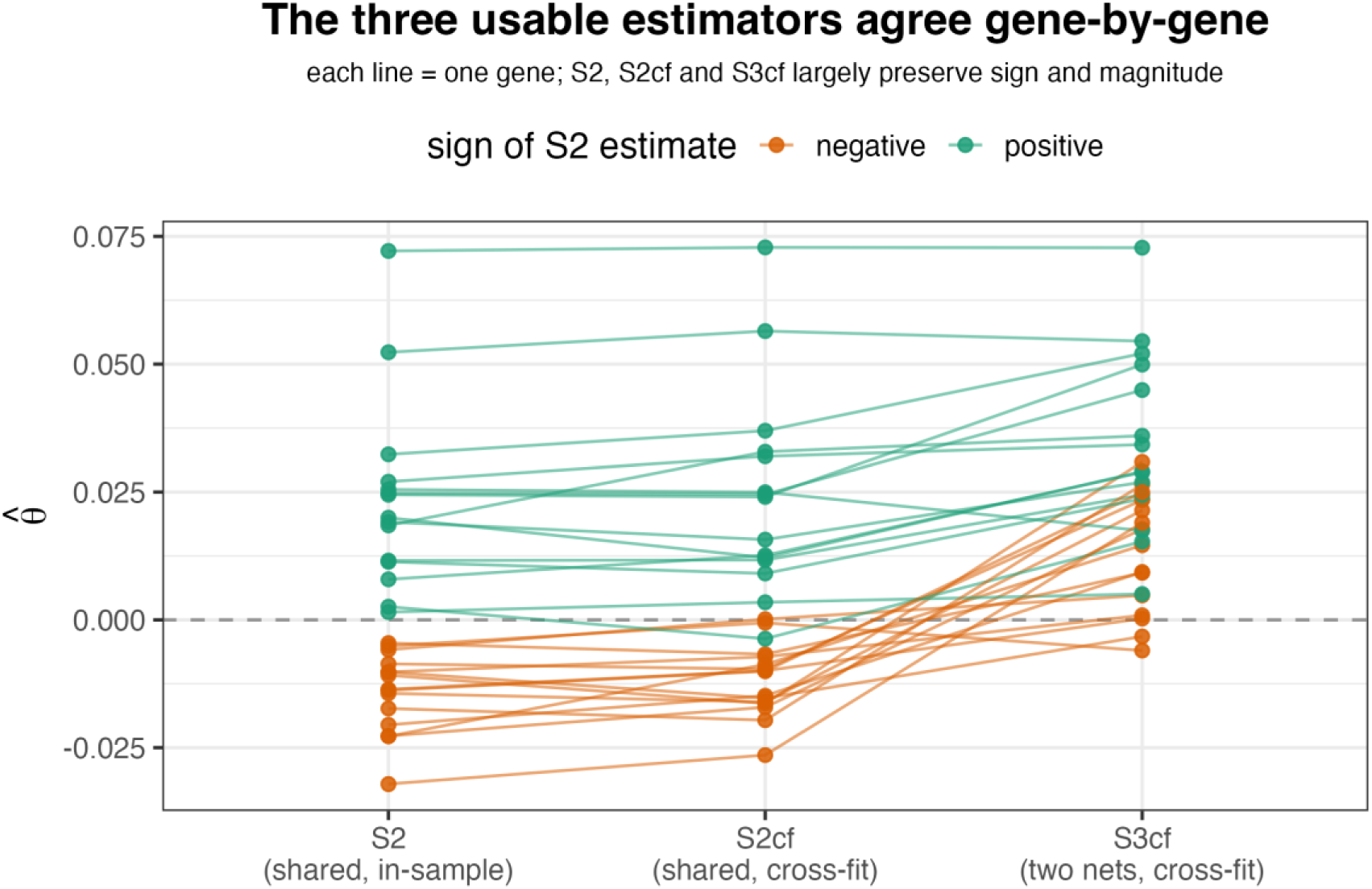
Parallel coordinate plot of gene-wise θ̂ for the three viable estimators (S2, S2cf, S3cf). Each line represents one gene, colored by the sign of the S2 estimate. All three show high consistency in both sign and magnitude.

### 3.6 Residual confounding diagnostic

The mean absolute correlation between the residuals and the background genes is low across all combinations (T_res about 0.012–0.024, Y_res about 0.014–0.022), with little difference. Note that the extremely low correlation of S1 in-sample’s T_res with X (1.2×10⁻⁴) is not “thorough deconfounding” but an artifact of overfitting crushing the residuals to numerical zero. This metric must therefore be interpreted cautiously for combinations with collapse, and is not used as a primary criterion in this paper.

## 4. Summary (at the results level)

Synthesizing the four classes of metrics across the six combinations yields a three-part pattern:

- **S1 (direct linear DML, no dimensionality reduction): fails in both in-sample and cross-fitting.**
- **S2 (shared unsupervised deep learning): optimal in-sample, and requires no cross-fitting.**
- **S3 (dual independent networks): collapses in-sample, and recovers under cross-fitting.**

This 2×3 pattern can be taken in at a glance (Figure 10): cross-fitting rescues S3, leaves S2 unchanged, and cannot rescue S1. The next two chapters give the theoretical explanation (Sec.5) and the verbal mechanistic reading, empirical corroboration, and method-selection advice (Sec.6)-readers interested only in the biological conclusions and methodological implications may go directly to Sec.6 and skip the derivations in Sec.5.

**Figure 10.**
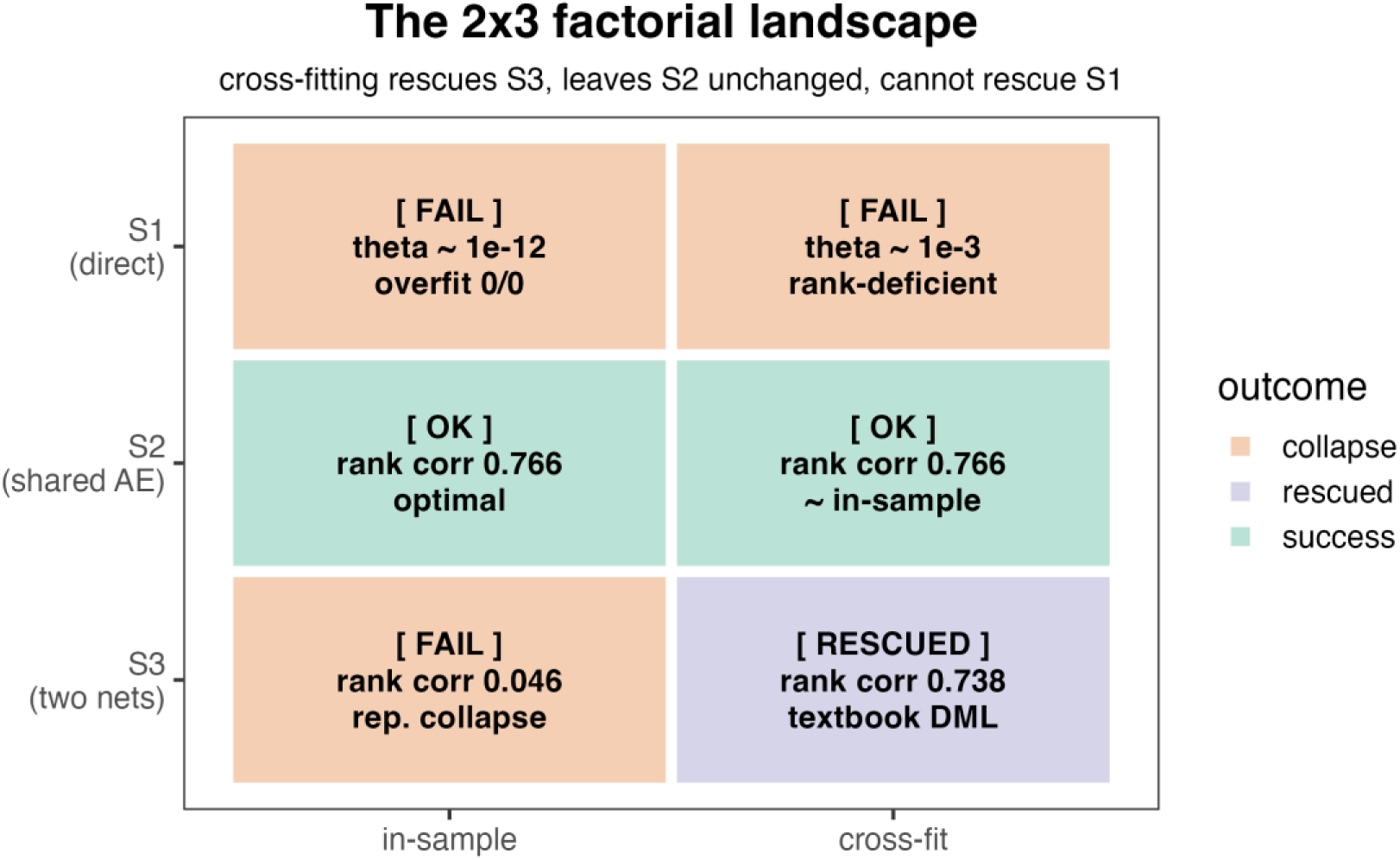
2×3 factorial landscape overview: cross-fitting rescues S3, does not change S2, cannot rescue S1.

## 5. Theoretical Foundations

This chapter derives, starting from the convergence rate condition of DML, the mathematical mechanism behind the three-part pattern. To facilitate selective reading, the dense formulas are concentrated here; readers interested only in the conclusions, mechanistic intuition, and empirical corroboration may go directly to Sec.6 Discussion-each subsection annotates the corresponding source in this chapter at the relevant place (e.g., “its mathematical mechanism is given in Sec.5.1”).

### 5.1 The rate condition of DML: the second-order product tail term and the passing threshold

#### Origin of the rate condition: the second-order product tail term

To understand why the “passing threshold” of the product rate condition in Sec.1.1 is *n*^−1/4^, we must return to the error expansion of *θ̂*. Denote the estimation errors of the two nuisance functions by Δ*l* = *l̂* − *l* and Δ*m* = *m̂* − *m*. Under the partially linear model *Y* = *θ*_0_*T* + *m*(*X*) + *ε* (where *ε* is the outcome noise, *E*[*ε*|*T*, *X*] = 0), denote

*T̃* = *T* − *l*(*X*) as the treatment residual. According to the expansion of Chernozhukov et al. (2018, Theorem 3.1), cross-fitting (sample splitting) makes the estimation error independent of the noise in the held-out fold, thereby **eliminating all first-order terms** (*E*[Δ*l* ⋅ *ε*] = *E*[Δ*m* ⋅ *T̃*] = 0) and leaving only a **second-order product term** after sample averaging:

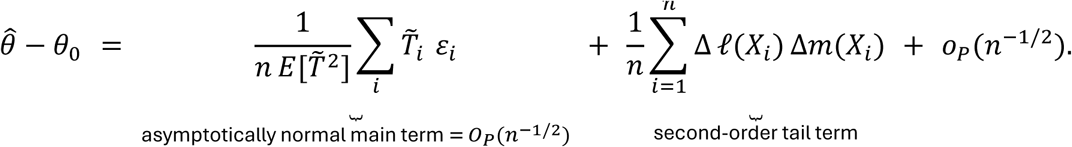

For this tail term not to interfere with 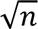 convergence, we need 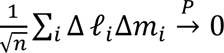. By Cauchy–Schwarz, 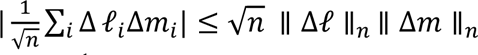 (where ∥⋅∥ is the empirical L2 norm, 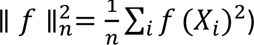, so a sufficient condition is precisely the product rate

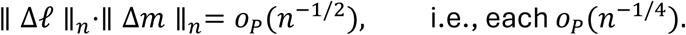

Under mild conditions the empirical L2 norm is asymptotically equivalent to the population L2 norm ∥⋅∥_2_; to facilitate the connection with Stone’s (1982) nonparametric rate results, we state everything below in terms of ∥⋅∥_2_, which the reader may understand as the probability-limit counterpart of the empirical L2 norm.

**This tail term directly translates “the bias of the causal estimate” into “the product of the fitting accuracies of the two nuisance functions”**: the finite-sample bias of *θ̂* depends only on how far *l̂* is from *l* and how far *m̂* is from *m*. The orthogonal score removes the first-order impact, cross-fitting severs the correlation between the estimation error and the noise, and what is left is purely “fitting residual”-the stronger the approximation ability of the nuisance functions, the smaller the tail term and the more accurate *θ̂*. (An intuitive reading of this mechanism is given in Sec.6.1.)

#### How high is the passing threshold: a nonparametric-rate perspective

DML’s rate condition, decomposed onto each nuisance function, requires each to individually reach *o*_*P*_(*n*^−1/4^)-note that this is a requirement on the **L2 norm** ∥ *m̂* − *m* ∥_2_, equivalently a requirement that the **mean squared error** ∥ *m̂* − *m* ∥^2^= *o*_*P*_(*n*^−1/2^). For an *s*-times differentiable function on ℝ^*d*^, the optimal nonparametric MSE convergence rate is *n*^−2*s*/(2*s*+*d*)^, i.e., an L2-norm rate of *n*^−*s*/(2*s*+*d*)^ ^[22]^. For the latter to reach *o*(*n*^−1/4^), we need

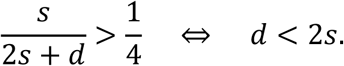

(Equivalently, requiring the MSE rate *n*^−2*s*/(2*s*+*d*)^ to reach *o*(*n*^−1/2^) likewise yields *d* < 2*s*.)

This threshold is more **stringent** than intuition suggests: for example, a twice-differentiable function (*s* = 2) requires *d* < 4, and a Lipschitz function (*s* = 1) requires *d* < 2-in the original feature dimension (p≈2000 in this study, p≈2×10⁴ for the full transcriptome) this is simply impossible to satisfy. Therefore **whether the threshold can be passed depends entirely on whether the true function possesses an exploitable low-dimensional effective structure** (sparsity, additive structure, low-dimensional manifold, compositional structure) that makes the effective dimension *d*^∗^ far smaller than the input feature count *p* and brings it below 2*s*. Different methods rely on different structural assumptions: Lasso relies on sparsity, random forests on low-dimensional additive structure, and deep neural networks on compositional structure ^[23]; [24]^. DML’s rate condition is “means-neutral” with respect to the nuisance-function estimator-**any model that can adapt to a low-dimensional effective structure (***d*^∗^ < 2*s***) can pass; whereas a model that only imposes a non-dimensionality-reducing structural assumption (e.g., a linear/parametric model on the raw high dimension), or fails to compress the effective dimension below** 2*s***, cannot pass.** But in a concrete application, which structural assumption holds and which method therefore qualifies is a question that must be analyzed in light of the data characteristics (see Sec.5.2).

#### Bias–variance decomposition of the rate condition

The ∥ *m̂* − *m* ∥_2_ in the rate condition ∥ *m̂* − *m* ∥_2_= *o*_*P*_(*n*^−1/4^) is not an indivisible whole but can be classically decomposed into approximation bias and estimation variance. Taking expectation over the randomness of the training set *D* and denoting *m̄* (*X*) = *E*_*D*_[*m̂*(*X*)] as the average estimator, we have

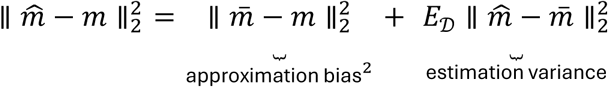

where the **approximation bias** ∥ *m̄* − *m* ∥ reflects whether the function class can express the true function: a non-decaying constant when a parametric model is misspecified, or decaying at the Stone rate above for a nonparametric model; the **estimation variance** reflects whether the complexity and regularization are appropriate: exploding for a high-capacity unregularized model, or *O*_*p*_(*n*^−1/2^) for a well-posed low-dimensional one. Because 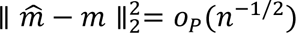 requires **both** terms to be individually *o*_*P*_(*n*^−1/2^), the rate condition is equivalent to two independent passing thresholds-approximation bias = *o*_*P*_(*n*^−1/4^) **and** estimation variance = *o*_*P*_(*n*^−1/2^); failure of either renders the whole unqualified. The two failure modes of single-cell DML each correspond to one component: **overfitting (variance-dominated)** means the estimation variance does not decay (the residuals are interpolated to zero and *θ̂* degenerates to 0/0); **underfitting (bias-dominated)** means the approximation bias does not decay (linear misspecification bias, or the curse of dimensionality at *p* ≈ 2000). This decomposition provides a unified anchor for Sec.5.2–Sec.5.4: the success or failure of each of the three methods can be attributed to which of these two thresholds (or both) is breached.

Therefore, the watershed for passing the rate condition lies in **whether a low-dimensional effective structure can be exploited**: nonparametric/semiparametric methods (Lasso/RF/deep networks) must compress the effective dimension below 2*s* (the Stone threshold) and keep the estimation variance controlled; parametric models (such as linear) face a more fundamental predicament-misspecification of the function class makes the approximation bias a constant, independent of 2*s*. This is precisely the root cause of the failure of S1’s unregularized high-dimensional linear regression in both modes, discussed in the next section.

### 5.2 The structural failure of S1 and the dimensionality predicament of classical methods

S1 represents the most naive class of approach: fitting the nuisance functions directly on the raw high-dimensional background X with **unregularized linear regression**-apart from the linear parametric assumption, it imposes no structural assumption (regularization, dimensionality reduction, sparsity, or composition) that **could effectively lower the effective dimension**. It should be clarified that S1 does not “have no structural assumption” but rather **imposes only one structural assumption that neither reduces dimensionality nor accords with the data-generating process** (linear + parametric)-an “assumption present but ineffective” approach. This section argues from two levels that such methods are infeasible in single-cell DML: first, the fixed parameter space of a parametric model keeps the approximation bias from decaying with sample size, and high-dimensional unregularized fitting makes the estimation variance explode; second, even classical nonparametric methods suffer the curse of dimensionality under high dimensionality (p≈2000).

#### Systematic failure of parametric models

The unregularized linear regression used by S1 is a parametric model: its parameter-space dimension is fixed at *p* (the number of background genes) and does not expand with sample size *n*. The defining feature of a parametric model is that the function class is fully determined by finitely many parameters-once the true function lies outside this class (model misspecification), the approximation bias is a non-decaying constant: ∥ *m̂* − *m* ∥_2_→ *c* > 0 (*n* → ∞). Single-cell gene regulation has a highly nonlinear biological basis-transcription-factor cooperativity, signaling-pathway cascades, feedback loops, epigenetic modifications, etc.-so the true *E*[*T*|*X*] and *E*[*Y*|*X*] are unlikely to be linear functions of X. Linear regression, as a special case of a parametric model, has a fixed function class (linear combinations of X); its approximation bias is structural and irreducible, so the rate condition ∥ *m̂* − *m* ∥_2_= *o*_*P*_(*n*^−1/4^) cannot be satisfied in the first place. In our experiments, the two modes of S1 embody this systematic failure in different forms:

- **In-sample mode = overfitting (variance-dominated)**: *p* = 1999 ≈ *n* = 2120, and unregularized linear regression can nearly interpolate the training data. As *p*/*n* → *γ*, the prediction error of linear regression converges to a nonzero constant related to *γ*, so ∥ *m̂* − *m* ∥ does not tend to zero at all. The residuals are artificially compressed to near zero, the orthogonal score degenerates to 0/0, and *θ̂* is of order ∼10⁻¹² (numerical zero). This is model capacity being **excessive** relative to the sample size, not insufficient.
- **Cross-fitting mode = rank deficiency + approximation bias**: the per-fold training sample drops to *n* ≈ 1696 < *p* = 1999, the design matrix is rank-deficient, and the linear coefficients are non-unique; what cross-fitting removes is only the overfitting (variance), not the approximation bias of the parametric model (the constant from linear misspecification)-cross-fitting cannot rescue the two structural problems of “inadequate function-class approximation + rank deficiency.”

#### The curse of dimensionality for classical nonparametric methods

A natural follow-up: since parametric models (such as linear regression) do not work, can classical nonparametric methods (such as kernel regression, smoothing splines) pass the rate condition? The essential difference between nonparametric methods and parametric models is that their function class does not presuppose a fixed parametric form but is progressively refined as the sample size grows-theoretically capable of approximating any smooth function. But the price of this flexibility is the curse of dimensionality. According to the Stone nonparametric rate analysis in Sec.5.1, DML requires *d* < 2*s*. **Stone (1982) proved an upper bound: for an** *s***-times differentiable function on a** *d***-dimensional space, the fastest L2 convergence rate achievable by any nonparametric regression method that does not exploit additional structure is** *n*^−*s*/(2*s*+*d*)^**-no change of algorithm can beat it. For this rate to reach the DML threshold** *o*(*n*^−1/4^)**, we must have** *s*/(2*s* + *d*) > 1/4**, i.e.,** *d* < 2*s***. Plugging in concrete numbers: even a twice-differentiable true function (***s* = 2**) allows only** *d* < 4**, and the Lipschitz case (***s* = 1**) is even more stringent at** *d* < 2. In the *p* ≈ 2000-dimensional raw feature space, even with smoothness *s* = 2 one would need an effective dimension *d* < 4-impossible to satisfy for any nonparametric method operating directly on the raw 2000-dimensional space. The convergence rate of classical nonparametric methods such as kernel regression and smoothing splines strictly depends on the input dimension *d*; in high-dimensional space they require exponentially growing sample sizes to maintain a given estimation precision (the classic statement of the “curse of dimensionality”), so the convergence rate of ∥ *m̂* − *m* ∥_2_ is far slower than *n*^−1/4^and the rate condition cannot be met.

#### Technological progress only exacerbates this predicament

It is worth emphasizing that *p* is not a stable current parameter but a quantity that **keeps growing with sequencing technology**: early Smart-seq and 10x v1 offered about 2,000 usable genes, 10x v3, Parse Bio, and other new-generation technologies stably detect 3,000–5,000, single-cell multi-omics (scRNA+scATAC+protein) pushes the feature dimension to 10^4^–10^5^, spatial transcriptomics expands from hundreds of target genes to thousands, and unfiltered whole-transcriptome analysis directly gives *p* ≈ 2×10^4^. **The larger** *p* **is, the harder the Stone threshold** *d* < 2*s* **is to satisfy, and the more structural and irreversible the failure of purely classical nonparametric methods becomes.** Conversely, this also means that deep-learning methods such as S2 and S3cf, which actively exploit low-dimensional intrinsic structure through dimensionality-reducing representations or compositional structure, hold an advantage that is not a temporary present-day gain but will keep amplifying with advances in sequencing technology-they are the only method family with scalability for single-cell DML in the face of an ever-expanding feature space.

#### Bias–variance extrapolation to n≫p

This paper takes a p≈n small sample as the experimental testbed; the conclusions can be extrapolated to n≫p (the common scenario for large atlases) on the basis of the monotone trend of the Sec.5.1 bias–variance decomposition as *n* grows. Increasing *n* has an asymmetric effect on the two thresholds: the estimation variance decays with *n* (the variance side improves), whereas the approximation bias of a parametric model is a constant that does not decay with *n* (the bias side does not improve). Hence under n≫p, S1’s failure flips from “in-sample variance-dominated overfitting” to “bias-dominated underfitting”-the linear-misspecification bias is laid bare, the variance is no longer the bottleneck, and cross-fitting, even after removing the overfitting bias, cannot rescue the constant approximation bias on the bias side. By contrast, for S2 and S3cf: increasing *n* simultaneously improves the approximation quality of the AE representation and the deep networks (bias side) and tightens the estimation variance of the linear head and after cross-fitting (variance side); both thresholds become easier to satisfy under n≫p. This extrapolation does not rely on any prior assumption such as “the linear condition holds”; it is a direct corollary of the Sec.5.1 rate-condition framework as *n* grows.

In summary, against the two thresholds of Sec.5.1, S1 fails on both the bias and the variance sides: on the bias side, the linear function class misspecifies the nonlinear gene regulation, so the approximation bias ∥ *m̄* − *m* ∥_2_→ *c* > 0 does not decay; on the variance side, high-dimensional unregularized fitting makes the estimation variance explode (in-sample interpolation, cross-fitting rank deficiency). Its **sole imposed structural assumption (linear + parametric) neither reduces dimensionality nor accords with the true generating process**-it reaps no dimensionality-reduction dividend, bears the misspecification bias, and additionally loses variance control due to excess capacity. The next two sections, Sec.5.3 and Sec.5.4, argue how S2 and S3cf each cross this threshold through their own effective structural assumptions.

### 5.3 S2 requires no cross-fitting: a structural property

S2 uses a two-step construction of **unsupervised representation learning + low-dimensional linear regression**: the autoencoder compresses the high-dimensional background X (p≈2000) into a low-dimensional representation Z (d=32), and the nuisance functions for treatment and outcome are fitted by linear regression on Z. This construction brings a key property-”requiring no cross-fitting” is a structural property of S2, arising because its nuisance functions (the low-dimensional linear heads) belong to the Donsker class. We argue this structural property below; the applicability boundary of S2 relative to S3cf (Donsker class vs. non-Donsker class) is compared together in Sec.5.4.

The reason classical DML needs cross-fitting is quite specific: when a high-capacity supervised model (trees, deep networks, etc.) fits both *E*[*T*|*X*] and *E*[*Y*|*X*] on the full data at once, it “records” the individual noise of T and Y into the model, producing a self-projection overfitting bias of *O*_*p*_(*n*^−1/2^)-a bias that does not vanish under 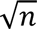 scaling and must be cut off by sample splitting (train within folds, predict on held-out folds). S2’s construction eliminates the source of this bias at two levels.

#### Level one: unsupervised representation learning blocks label leakage

S2’s autoencoder is trained only on X with the reconstruction objective ∥ *X* − *X̂* ∥^2^ and **never touches T and Y from start to finish**. Hence, with respect to T and Y, the representation *Ẑ* = *f̂*(*X*) is strictly exogenous-the AE cannot “remember” any noise pattern related to T or Y. Even if the AE overfits on X (e.g., remembers some individual noise of X), these overfitting artifacts are orthogonal to the structural residuals *η* = *T* − *E*[*T*|*X*] and *ε* = *Y* − *E*[*Y*|*X*] of T and Y-**at most introducing irrelevant noise dimensions into Z, adding variance but not bias**. This exogeneity isolation makes the representation-learning stage of S2 naturally immune to “label overfitting”-precisely the primary target that cross-fitting is meant to correct.

#### Level two: the low-dimensional linear head belongs to the Donsker class, with self-projection bias

*O*_*p*_(*n*^−1^). When doing linear regression on Z, *m̂* and *l̂* are linear projections in a 32-dimensional space, belonging to a low-complexity (Donsker) parametric class. The Neyman orthogonal score removes the first-order bias, leaving a residual bias in product form ∥ *m̂* − *m* ∥⋅∥ *l̂* − *l* ∥ (Sec.5.1). Under the well-posed conditions of OLS at d=32, n=2120, the estimation error is naturally at the parametric rate *O*_*p*_(*n*^−1/2^)-this is a purely statistical property of OLS; regardless of whether the true *E*[*T*|*Z*] is linear, the rate at which the OLS coefficients converge to the best linear projection is *O*_*p*_(*n*^−1/2^). The product of the two -> *O*_*p*_(*n*^−1^), which tends to zero under 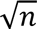 scaling. According to Chernozhukov et al. (2018), when the nuisance functions belong to the Donsker class, cross-fitting is not necessary and full-sample plug-in estimation suffices for 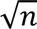-consistency.

The two levels combine: S2 eliminates from the root the bias that cross-fitting is meant to correct-the AE does not touch the labels -> representation learning has no label overfitting; the low-dimensional linear head -> the self-projection bias in the regression stage is *O*_*p*_(*n*^−1^) -> no sample splitting is needed. Against the two thresholds of Sec.5.1: on the variance side, the estimation error of the low-dimensional Donsker-class linear head is at the parametric rate *O*_*p*_(*n*^−1/2^) (estimation variance *O*_*p*_(*n*^−1^), satisfying *o*_*P*_(*n*^−1/2^)), passing; the self-projection bias

*O*_*p*_(*n*^−1^) vanishes under 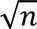 scaling and poses no burden on the bias side. **Note that “no cross-fitting required” is a structural property of S2, independent of the functional form of** *E*[*T*|*Z*] **or** *E*[*Y*|*Z*]**-but it only guarantees the rate of convergence toward the best linear projection, not that the linear projection itself is free of approximation bias; the latter depends on whether the AE representation Z is a sufficient dimensionality reduction, and is where the applicability boundary of S2 relative to S3cf lies (see Sec.6.4 for details).**

### 5.4 S3cf is effective: the three-layer chain of standard DML

S3cf falls squarely within the standard framework of Chernozhukov et al. (2018): use two (sufficiently approximating) supervised learners to estimate *E*[*T*|*X*] and *E*[*Y*|*X*] separately, eliminate the overfitting bias with cross-fitting, and then estimate θ with the Neyman orthogonal score. Unlike S2’s linear head, S3cf’s supervised networks fit the conditional expectations on the raw high-dimensional X end-to-end, belonging to a high-capacity non-Donsker class that requires cross-fitting to eliminate the self-projection bias-this is precisely where the standard DML recipe is broad-spectrum for high-dimensional causal inference. We now give the complete chain by which it satisfies the DML theoretical framework.

#### Layer one: the score function and Neyman orthogonality

S3cf uses the orthogonal score

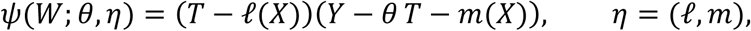

where the nuisance functions *l*(*X*) = *E*[*T*|*X*] and *m*(*X*) = *E*[*Y*|*X*] are directly the **conditional expectations on the raw high-dimensional X**, fitted end-to-end by two independent deep networks-without any low-dimensional projection. At the true value this score satisfies the Neyman orthogonality condition *δ*_*η*_*E*[*Ψ*(*W*; *θ*_0_, *η*_0_)][*η* − *η*_0_] = 0, so the first-order estimation error of the nuisance functions is removed, leaving only the second-order product tail term 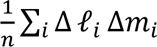 of Sec.5.1. This step holds for **any** nuisance-function estimator, regardless of whether the function is linear or dimensionality-reducing.

#### Layer two: the deep networks satisfy the rate condition

The crux is whether the two supervised networks can make ∥ Δ*l* ∥_2_and ∥ Δ*m* ∥_2_each reach *o*_*P*_(*n*^−1/4^). Deep nonparametric regression theory provides a directional basis: **if** the true *E*[*T*|*X*] and *E*[*Y*|*X*] possess **compositional / hierarchical composition structure** (generalized hierarchical composition) and the network architecture matches the compositional structure of the true function, then deep networks with ReLU activation can circumvent the curse of dimensionality and achieve convergence near the parametric rate *Õ*_*P*_(*n*^−*β*/(2*β*+*d*)^) (where *d*^∗^ is the effective intrinsic dimension rather than the raw dimension *p*; Schmidt-Hieber [25]; [26]). As long as this rate is faster than *n*^−1/4^-which holds when the effective dimension *d*^∗^is moderate and the smoothness *β* is sufficient-the product error ∥ Δ*l* ∥_2_∥ Δ*m* ∥_2_= *o*_*P*_(*n*^−1/2^) and the rate condition is satisfied. Note: the above theoretical result requires the network structure to be specially designed for the compositional structure of the true function, whereas the fixed 8-layer fully connected architecture in this study is a heuristic implementation of that theory and does not strictly meet its premises. **The high approximation capacity of deep networks is precisely what drives this second-order tail term below** 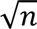.

#### Layer three: cross-fitting eliminates the self-projection bias (indispensable)

Unlike S2’s low-dimensional linear head, S3’s supervised networks are a **high-capacity non-Donsker class**: when trained on the full data with (*X*, *T*) and (*X*, *Y*) at once, the network “memorizes” the individual noise of *T* and *Y*, producing a self-projection overfitting bias of *O*_*P*_(*n*^−1/2^)-a bias that does not vanish under 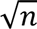 scaling and would break the asymptotic normality. Cross-fitting (train within folds, predict on held-out folds) makes the nuisance-function estimate independent of the noise of the sample being evaluated, thereby removing this bias. **This is the fundamental difference why S3 must cross-fit while S2 need not**: S3’s nuisance functions are fitted directly under supervision on the labels and belong to a high-capacity class; S2’s representation is unsupervised (does not touch labels) and its linear head belongs to a low-dimensional Donsker class (Sec.5.3). Clarification: cross-fitting removes the **self-projection bias on the nuisance-function bias side**, not the variance; once it allows the second layer’s rate condition to be realized, the variance of *θ̂* is governed by the asymptotically normal main term of Sec.5.1, 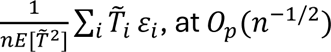-this is precisely the 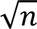-consistency of DML, and it also echoes the S3cf intra-gene variance measured in Sec.3.4 (4.67×10⁻⁴, the same order as S2).

#### Theoretical implication of broad-spectrum applicability

The validity of S3cf does not depend on a conditional-expectation assumption that the nuisance functions belong to the Donsker class-it imposes no “belongs to the Donsker class” functional constraint on the modeling of *E*[*T*|*X*] and *E*[*Y*|*X*]: as long as the deep networks can approximate them (layer two) and cross-fitting removes the bias (layer three), S3cf gives a 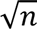-consistent estimate. This is where it is “broad-spectrum” relative to S2; S3cf trades an end-to-end network x cross-fitting for “theoretical robustness without this assumption.” Hence the effectiveness of S3cf needs no new theory-it is simply the correct instantiation of standard DML in the single-cell setting.

## 6. Discussion

This chapter reads in words how the theory of Sec.5 is borne out in the data, and gives method-selection advice. To facilitate selective reading, formulas are not repeated here; for the rigorous derivation one may return to the parenthetically noted corresponding section (e.g., “see Sec.5.1”).

### 6.1 Why S1 fails in both modes-and the dimensionality predicament of classical methods

**Mechanistic reading.** The two failure modes of S1 both appear to point to “parametric models don’t work,” but the **mechanisms are entirely different, and neither can be simply attributed to “parametric-model capacity is too low.”** The essence of in-sample failure is **overfitting**: in the critical regime of p≈n, unregularized linear regression can nearly interpolate the training data, artificially compressing the residuals to near zero, and the orthogonal score degenerates to 0/0-this is model capacity being **excessive** relative to the sample size, not insufficient. The essence of cross-fitting failure is **rank deficiency + approximation bias**: the per-fold sample drops to 1696, below the background dimension 1999, so the coefficients are non-unique; at the same time the parametric model has an irreducible approximation error for the true nonlinear confounding structure. Cross-fitting, while removing the overfitting bias, cannot rescue the two structural problems of “inadequate function-class approximation + rank deficiency” (mathematical mechanism in Sec.5.1–5.2).

**Why can cross-fitting rescue S3 but not S1?** One-sentence summary: cross-fitting treats the one disease of “a high-capacity model memorizing label noise”; S1 suffers from a different disease-”function-class capacity inadequacy + training-fold rank deficiency”-**the wrong medicine for the disease.** (Technical mechanism in Sec.5.1–5.2.)

**It is also foreseeable in theory**: even if the sample size were raised to the millions, **imposing only an ineffective structural assumption (a non-dimensionality-reducing parametric model)** would merely flip “overfitting” into “underfitting with non-decaying bias,” still unable to pass the threshold of the rate condition (the asymmetric trend of bias vs. variance as *n* grows is in the Sec.5.2 extrapolation).

**What it needs is not “more data” but “better structural exploitation.”**

**Why classical nonparametric methods may not save the day either.** A natural follow-up: can classical nonparametric methods (such as kernel regression, smoothing splines) qualify? Sec.5.2 gives a negative answer from the curse-of-dimensionality angle. In other words, merely “switching to another nonparametric function class” without solving the dimensionality problem is likewise a dead end in single-cell DML.

**Can classical models that depend on a structural assumption succeed? The key lies in whether the structural assumption holds**, and the structural assumptions relied upon by different methods **do not align equally with the characteristics of single-cell data.**

**Lasso relies on a sparsity assumption**: only when *s* ≪ *p* genes truly drive T or Y can Lasso achieve the “dimensionality-escaping” rate 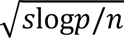. But gene-expression regulation is **highly polygenic**-the polygenicity evidence from GWAS, the omnigenic-model hypothesis, and the widespread weak-correlation structure in single-cell transcriptomes all point to the picture of “hundreds to thousands of genes collectively participating in regulation through weak signals.” Under this picture Lasso’s sparsity assumption **does not hold naturally**; forced top-k truncation would omit most weak signals and the rate condition would be breached.

**Random forests rely on low-dimensional effective sparse structure**: analyses such as Scornet’s show that RF can circumvent the curse of dimensionality under structural assumptions like “sparse additive,” but under general high-dimensional dense signals RF’s convergence rate is still plagued by the curse of dimensionality.

Single-cell nonlinearity comes from transcription-factor cooperativity, pathway cascades, feedback loops, and other **distributed non-additive interactions**, which do not match a “few variables + additive” structure.

**Deep networks rely on a low-dimensional manifold or compositional/hierarchical structure** ([27], [28]): these two classes of assumption **are precisely the features repeatedly empirically supported for single-cell data**-low-dimensional manifold structure is widely validated by scVI, UMAP, diffusion maps, etc., and the compositional structure of gene regulation naturally matches the hierarchical composition of deep networks.

In other words, although Lasso (regularized parametric model), RF (nonparametric ensemble), and deep networks can all “circumvent the curse of dimensionality through a structural assumption” **in form**, the **degree of alignment** between the structural assumption each requires and the empirically demonstrated characteristics of single-cell data differs substantively-Lasso and RF require a sparsity or additive structure that single-cell data do not possess, whereas deep networks require the low-dimensional manifold and compositional structure that single-cell data have been repeatedly validated to possess. Therefore the conclusion of this paper is not merely “S1’s structural assumption is ineffective -> failure,” but further: **in the single-cell context, deep learning is currently the only method family whose structural assumption truly aligns with the data characteristics. The bias–variance decomposition (Sec.5.1) pinpoints this divide: Lasso and RF could pass on the variance side (regularization/ensemble) but fail on the bias side-structural-assumption mismatch keeps the approximation bias from decaying; by contrast, S1 fails on both the bias and variance sides, while S2/S3cf pass on both. This distinction is invisible from the overall rate condition alone (all three can satisfy the rate when their assumptions hold); only the bias–variance decomposition reveals that Lasso/RF fail on the bias side.** Lasso, random forests, and other classical methods were not included in our experiments, but even if they were, there is a substantive risk in theory that their structural assumptions do not match the polygenic / distributed non-additive regulatory characteristics of single-cell data and thus cannot attain the rate condition-this is an open question that this paper does not answer experimentally (see Sec.7 Limitations).

### 6.2 Why S2 requires no cross-fitting-intuitive reading and empirical corroboration

**Why it “does not need to be rescued.”** Cross-fitting fixes one very specific thing: a high-capacity supervised model that both learns the nuisance functions on the same batch of samples and is then used to compute the residuals will “record” the sample noise into the model, producing a layer of overfitting bias. But S2’s representation Z is learned **unsupervised**-the autoencoder does only one thing: compress X and reconstruct it, and the objective function **never looks at T and Y from start to finish**. Since representation learning never touches the labels, “recording label noise into the model” simply cannot happen; moreover, Z has only 32 dimensions, far smaller than the sample size, so linear regression on it is well-posed and not overfit. Thus the layer of bias that cross-fitting is meant to remove is essentially absent from the source in S2 (rigorous derivation in Sec.5.3, exogeneity isolation and the low-dimensional Donsker-class argument).

There is also a layer of “structural insurance”: because the representation is determined solely by X and is by definition uncorrelated with the T/Y residuals, even if the autoencoder slightly overfits on X, that bit of “memory” only adds to Z some noise dimensions irrelevant to the causal estimate-**adding variance, not bias** (Sec.5.3 exogeneity isolation).

**Empirical demonstration.** This theoretical expectation is strongly supported by the data: S2 and S2cf are nearly identical across all metrics-resolution 5.60 vs 5.59×10⁻⁴, biological rank correlation 0.766 vs 0.766, sign consistency 0.933, shrinkage factor β = 0.975. Cross-fitting brings essentially no change to S2, exactly as the theory predicts. This high consistency does not “prove” that S2 needs no cross-fitting (that conclusion is given directly by the structural argument of Sec.5.3) but provides an intuitive empirical demonstration for the theory: if S2’s construction truly eliminates from the root the overfitting bias that cross-fitting is meant to correct, then adding cross-fitting should make no difference-and that is exactly what the experiment shows. Of course, requiring no cross-fitting only guarantees the rate of convergence toward the best linear projection, not that the linear projection itself is free of approximation bias; the boundary of the latter is in Sec.6.4.

### 6.3 Why cross-fitting is indispensable for S3-intuitive reading and empirical corroboration

**Why it “must be rescued.”** S3 is the exact counterpart of S2: it uses two **supervised** networks to fit *E*[*T*|*X*] and *E*[*Y*|*X*] end-to-end, with high capacity and learning directly from the labels. When trained on the full data, the networks record the individual noise of T and Y, producing a layer of overfitting bias that does not vanish at the 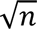 scale, enough to break the asymptotic normality of the estimate. Cross-fitting (train within folds, predict on held-out folds) makes “the samples used to learn the nuisance functions” and “the samples used to compute the residuals” mutually independent, severing this layer of bias (note that cross-fitting removes this bias, not the variance; the variance of θ̂ is determined by the asymptotic main term, see Sec.5.4). This is the fundamental difference why S3 must cross-fit while S2 need not: **S3’s nuisance functions are fitted directly under supervision on the labels and belong to a high-capacity class; S2’s representation is unsupervised and its linear head belongs to a low-dimensional Donsker class** (rigorous chain in Sec.5.4).

**Empirical corroboration.** This reasoning is precisely borne out by the results: S3 in-sample collapses (rank correlation only 0.046); S3cf is rescued (rank correlation 0.738, resolution improved by about six orders of magnitude)-remove cross-fitting and the high-capacity supervised network immediately overfits and collapses; restore cross-fitting and θ̂ recovers to a level comparable to S2. Sec.3.4 further shows that the stability of S3cf (intra-gene variance 4.67×10⁻⁴) is of the same order as S2 (2.7×10⁻⁴)-”genuine stability backed by a real signal,” fundamentally different from S3 in-sample’s “stably estimating a near-zero wrong value” (the 1.3×10⁻¹⁰ false stability).

**The meaning of broad-spectrum applicability.** The validity of S3cf does not depend on a conditional-expectation assumption that the nuisance functions belong to the Donsker class: as long as the deep networks can approximate them and cross-fitting removes the bias, S3cf gives reliable estimates. This makes it the “broad-spectrum” instantiation of standard DML in the single-cell setting-**”broad-spectrum” refers to the breadth of the theoretical scope of applicability (independence from the Donsker-class condition).** Here the object of comparison must be anchored at the **nuisance function**: S2’s nuisance functions are Donsker-class linear heads, whereas S3cf’s are non-Donsker-class deep networks; the two share the same backbone deep network only as a dimensionality-reduction/representation tool, and the function classes to which the nuisance functions belong are not the same. S3cf trades non-Donsker-class nuisance functions + cross-fitting for broad-spectrum applicability without the Donsker-class assumption. The real cost is higher computational demand (detailed in the next section).

### 6.4 Method selection: computational cost vs. broad-spectrum applicability

S2 and S3cf are both “effective” solutions, but suit different scenarios:

- **S2 (shared unsupervised deep learning, in-sample)-computationally economical, suited to large-scale screening.** S2 attains estimation quality comparable to standard cross-fitted DML at about 1/10 the computational cost.
- **S3cf (dual networks + cross-fitting)-the standard DML recipe, providing a broad-spectrum scheme when the conditional expectation may exceed the approximable range of the Donsker class.**

**Note that the broad spectrum of S3cf is still bounded by the approximation capacity of its non-Donsker-class deep networks-if the true** *E*[*Y*|*X*] **lies beyond the approximable range of the deep-network family, S3cf also fails (S2, whose nuisance functions are only Donsker-class linear heads, fails even earlier); at that point the problem is no longer at the method-selection level but at the technical boundary of the entire causal-inference tool family.**

In one sentence: **”cross-fitting can rescue S3, but S2 does not need to be rescued in the first place”**; in practice, trade off by “throughput first -> choose S2, broad-spectrum first -> choose S3cf.”

## 7. Limitations

1. **Regularized / classical nonparametric methods were not tested.** S1 represents only unregularized direct linear DML; classical methods such as Lasso and random forests were not included in the experimental comparison.

This is a design choice of this paper-to keep the clean control between S1 and S2 (the only difference being “whether dimensionality reduction is applied”). At the theoretical level, this paper has argued in Sec.5.2 and Sec.6.1 for the **structural disadvantages of these classical methods in single-cell DML, but did not test them empirically**.

1. 2. **Limited number of bootstraps.** In-sample stability used N = 6 bootstraps, with low degrees of freedom (df = 5); the stability figures are “pilot”-level relative comparisons and should not be over-interpreted in absolute precision. **The stability test of S3cf is especially constrained**: due to the computational constraint (each estimate involves 2 deep networks x 5-fold training), the bootstrap of S3cf was completed with only N = 3 x 10 genes (vs. N = 6 x 30 genes for the three in-sample strategies); its stability figures are order-of-magnitude pilots rather than precise estimates.
2. **A single cell type, a single dataset.** Validation was performed only on memory B cells (2,120 cells) of GSE189050; generalizability across cell types and datasets remains to be tested.
3. **No ground truth.** Observational data lack true causal-effect values, so evaluation relies on indirect metrics such as resolution / biological plausibility / stability, rather than on the estimation error itself.

## 8. Conclusion

In single-cell transcriptomics, the success or failure of double-machine-learning causal inference depends on how the nuisance functions are constructed. Through a 3-strategy x (in-sample / cross-fitting) 2×3 factorial experiment, we obtain a three-part pattern with a clear theoretical explanation: direct linear DML (S1) fails in both modes because it structurally violates the rate condition; shared unsupervised deep learning (S2) naturally satisfies the DML requirements by virtue of “representation independent of labels + well-posed low-dimensional regression,” and is optimal in-sample with no need for cross-fitting; dual independent networks

(S3) are the standard DML recipe and require cross-fitting to take effect. Accordingly, we recommend: **for large-scale gene screening, prioritize S2 to save computation; when the conditional expectation may exceed the approximable range of Donsker-class nuisance functions, use S3cf as a broad-spectrum fallback.** The methodological contribution of this paper is to place, for the first time, nuisance-function construction strategies and cross-fitting within the same factorial framework, revealing their division of labor and complementarity in single-cell DML. A deeper methodological implication is that in large-scale single-cell causal inference, the standard DML recipe with the broadest theoretical assumptions (S3cf) and the most computationally economical shared-unsupervised-deep-learning method (S2) can form a synergistic partnership through cross-validation-the former provides broad-spectrum reliability endorsement on a feasible subset, while the latter achieves estimation at the whole-transcriptome scale. Although their nuisance functions belong to different function classes (S2 to the Donsker class, S3cf to the non-Donsker class), they share the same backbone deep network as a dimensionality-reduction tool; therefore, in scenarios where S3cf and S2 are highly consistent, one can confidently use S2 to complete the screening, providing a feasible path for large-scale causal screening under computational constraints.

## Notes

### Competing Interest Statement

The authors have declared no competing interest.

## REFERENCES

[1] Chernozhukov V, Chetverikov D, Demirer M, Duflo E, Hansen C, Newey W, Robins J. Double/debiased machine learning for treatment and structural parameters. The Econometrics Journal. 2018;21(1):C1–C68. doi:10.1111/ectj.12097

[2] Dong M, Wang B, Wei J, et al. Causal identification of single-cell experimental perturbation effects with CINEMA-OT. Nat Methods. 2023;20(11):1769–1779. doi:10.1038/s41592-023-02040-5

[3] An S, Cho JW, Cao K, Xiong J, Hemberg M, Wan L. scCausalVI disentangles single-cell perturbation responses with causality-aware generative model. Cell Syst. 2025;16(11):101443. doi:10.1016/j.cels.2025.101443

[4] Wang JC, Chen YJ, Zou Q. GRACE: Unveiling Gene Regulatory Networks With Causal Mechanistic Graph Neural Networks in Single-Cell RNA-Sequencing Data. IEEE Trans Neural Netw Learn Syst. 2025;36(5):9005–9017. doi:10.1109/TNNLS.2024.3412753

[5] Papili Gao N, Ud-Dean SMM, Gandrillon O, Gunawan R. SINCERITIES: inferring gene regulatory networks from time-stamped single cell transcriptional expression profiles. Bioinformatics. 2018;34(2):258–266. doi:10.1093/bioinformatics/btx575

[6] Dixit A, Parnas O, Li B, et al. Perturb-Seq: Dissecting Molecular Circuits with Scalable Single-Cell RNA Profiling of Pooled Genetic Screens. Cell. 2016;167(7):1853–1866.e17. doi:10.1016/j.cell.2016.11.038

[7] Ishikawa M, Sugino S, Masuda Y, et al. RENGE infers gene regulatory networks using time-series single-cell RNA-seq data with CRISPR perturbations. Commun Biol. 2023;6(1):1290. Published 2023 Dec 28. doi:10.1038/s42003-023-05594-4

[8] Qiu X, Rahimzamani A, Wang L, et al. Inferring Causal Gene Regulatory Networks from Coupled Single-Cell Expression Dynamics Using Scribe. Cell Syst. 2020;10(3):265–274.e11. doi:10.1016/j.cels.2020.02.003

[9] Tejada-Lapuerta A, Bertin P, Bauer S, Aliee H, Bengio Y, Theis FJ. Causal machine learning for single-cell genomics. Nat Genet. 2025;57(4):797–808. doi:10.1038/s41588-025-02124-2

[10] Wei Z, Wang Y, Gao Y, et al. Benchmarking algorithms for generalizable single-cell perturbation response prediction. Nat Methods. 2026;23(2):451–464. doi:10.1038/s41592-025-02980-0

[11] Chevalley M, Roohani YH, Mehrjou A, Leskovec J, Schwab P. A large-scale benchmark for network inference from single-cell perturbation data. Commun Biol. 2025;8(1):412. Published 2025 Mar 11. doi:10.1038/s42003-025-07764-y

[12] Farrell MH, Liang T, Misra S. Deep neural networks for estimation and inference. Econometrica. 2021;89(1):181–213. doi:10.3982/ECTA16901

[13] Du JH, Zeng Z, Kennedy EH, Wasserman L, Roeder K. Causal Inference for Genomic Data with Multiple Heterogeneous Outcomes. J Am Stat Assoc. Published online June 6, 2025. doi:10.1080/01621459.2025.2468014

[14] Lopez R, Regier J, Cole MB, Jordan MI, Yosef N. Deep generative modeling for single-cell transcriptomics. Nat Methods. 2018;15(12):1053–1058. doi:10.1038/s41592-018-0229-2

[15] Eraslan G, Simon LM, Mircea M, Mueller NS, Theis FJ. Single-cell RNA-seq denoising using a deep count autoencoder. Nat Commun. 2019;10(1):390. Published 2019 Jan 23. doi:10.1038/s41467-018-07931-2

[16] Wang FA, Yi C, Chen J, He R, Liu J, Li Y. scTFBridge: a disentangled deep generative model informed by TF-motif binding for gene regulation inference in single-cell multi-omics. Nat Commun. 2025;16(1):9166. Published 2025 Oct 15. doi:10.1038/s41467-025-64227-y

[17] Dimitrov D, Schrod S, Rohbeck M, Stegle O. Interpretation, extrapolation and perturbation of single cells. Nat Rev Genet. 2026;27(5):349–370. doi:10.1038/s41576-025-00920-4

[18] Zinati Y, Takiddeen A, Emad A. GRouNdGAN: GRN-guided simulation of single-cell RNA-seq data using causal generative adversarial networks. Nat Commun. 2024;15(1):4055. Published 2024 May 14. doi:10.1038/s41467-024-48516-6

[19] Wei H, Lu H, Zhao H. Inferring Time-Lagged Causality Using the Derivative of Single-Cell Expression. Int J Mol Sci. 2022;23(6):3348. Published 2022 Mar 20. doi:10.3390/ijms23063348

[20] Christensen OP, Markham A, Kang H, Gabriel E, Pers TH. Causal effect estimation from trans-regulatory single-cell CRISPR screens. Cell Genom. 2026;6(6):101251. doi:10.1016/j.xgen.2026.101251

[21] Slight-Webb S, Thomas K, Smith M, et al. Ancestry-based differences in the immune phenotype are associated with lupus activity. JCI Insight. 2023;8(16):e169584. Published 2023 Aug 22. doi:10.1172/jci.insight.169584

[22] Stone CJ. Optimal global rates of convergence for nonparametric regression. Ann Statist. 1982;10(4):1040–1053. doi:10.1214/aos/1176345969

[23] Schmidt-Hieber J. Nonparametric regression using deep neural networks with ReLU activation function. Ann Stat. 2020;48(4):1875–1897. doi:10.1214/19-AOS1875

[24] Bauer B, Kohler M. On deep learning as a remedy for the curse of dimensionality in nonparametric regression. Ann Stat. 2019;47(4):2261–2285. doi:10.1214/18-AOS1747

